# Clonal interference and changing selective pressures shape the escape of SARS-CoV-2 from hundreds of antibodies

**DOI:** 10.1101/2025.01.22.634135

**Authors:** Hugh K. Haddox, Omar Abdel Aziz, Jared G. Galloway, Javen Kent, Cameron R. Cooper, Jesse D. Bloom, Frederick A. Matsen

## Abstract

SARS-CoV-2 rapidly evolves to evade human immunity. While the virus’s overall resistance to human polyclonal antibody responses has steadily increased over time, the dynamics by which it escaped individual monoclonal antibodies within these responses have not been thoroughly explored. Recently, a series of studies by Cao et al. [1, 2, 3] used deep mutational scanning (DMS) to identify which mutations allow the Wuhan-Hu-1 receptor-binding domain to escape binding by individual antibodies, doing so for thousands of antibodies. Here, we sought to use these data to retrospectively examine the evolutionary dynamics of escape from a set of 1,603 antibodies. For each antibody, we used the DMS data to predict an antibody-escape score for each of thousands of globally circulating viral sequences from the first 3.5 years of the pandemic, and then computed an escape trajectory that quantifies how the population’s average escape score changed over time. We use pseudovirus neutralization data from Cao et al. and Wang et al. [4] to validate common patterns in escape trajectories. While some trajectories increase monotonically over time, others show large fluctuations as a result of clade-displacement events that reduce the frequency of antibody-escape mutations in the viral population. Fitness effects of mutations estimated from natural sequences suggest that the mutations are displaced due to clonal interference. Further, these estimates suggest that the order in which escape mutations arose is shaped by changing selective pressures. Overall, this work helps describe how SARS-CoV-2 evaded the individual components of a polyclonal immune response in nature, and suggests that evasion occurred via complex evolutionary dynamics.

## Introduction

SARS-CoV-2 rapidly evolves, in part to escape human antibody-mediated immunity. For instance, in the first few years of the pandemic, the field isolated hundreds of neutralizing monoclonal antibodies from humans [1, 2, 3, 4, 5, 6, 7, 8, 9]. These antibodies target multiple epitopes on the spike protein, and several antibodies were approved as therapeutics. As the virus began to evolve resistance to some antibodies, interest turned towards identifying ones with broad neutralizing activity against emerging viral variants, as well as cross-reactivity with other sarbecoviruses, with the hope that these antibodies might be less susceptible to viral escape [4, 10, 11, 12, 13, 14, 15]. However, within a matter of years, the virus evolved resistance to nearly all antibodies isolated within this initial period of the pandemic [3, 16]. As the pandemic continues and human antibody responses are updated upon new exposures, the virus continues to evolve resistance to these responses [17], making it important to understand the evolutionary processes at play.

How SARS-CoV-2 evolved resistance to human antibody-mediated immunity during the first few years of the pandemic is one of the best-documented stories of immune escape to date, and some aspects of the story are very clear. In general, the virus’s resistance to human sera and monoclonal antibodies tended to increase over time upon the appearance of new variants of concern [1, 2, 3, 13, 16, 18, 19, 20, 21, 22]. And key mutations that drove escape were identified using a combination of bioinformatics and laboratory experiments [1, 2, 3, 22, 23, 24, 25, 26, 27, 28, 29, 30, 31, 32, 33].

However, some aspects of this evolutionary story have not been fully resolved. For instance, while it is clear that the virus’s overall resistance to antibodies steadily increased, we still lack a complete story of how the virus evolved resistance to each of the large number of monoclonal antibodies that collectively made up the human polyclonal response. How did the virus navigate this complex immune fitness landscape in nature? Past studies have identified groups of antibodies that were escaped at different times by different mutations and variants of concern. However, these studies primarily focused on several well-studied antibodies, leaving room for a more comprehensive analysis.

Here, we sought to build on the above work by analyzing escape from hundreds of individual antibodies isolated from humans. Specifically, for each antibody, we sought to answer the following questions: When did the virus population escape the antibody? Did levels of escape in the viral population steadily increase over time, or were patterns more rugged? Which mutations drove escape? And how do patterns of escape relate to the frequent clade-displacement events in SARS-CoV-2’s evolution?

To answer these questions, we leveraged data from a series of studies by Cao et al. [1, 2, 3], which used high-throughput experiments to isolate and characterize thousands of human monoclonal antibodies from the first few years of the pandemic. Specifically, they isolated antibodies that bound to the SARS-CoV-2 spike receptor-binding domain (RBD), which is a major target of neutralizing antibodies [28, 34]. For each antibody, they then used deep mutational scanning (DMS) to identify which mutations allow the Wuhan-Hu-1 RBD to escape antibody binding. They also measured the neutralization potency of these antibodies against pseudoviruses bearing spike proteins of several variants of concern, finding that the virus evolved resistance to nearly all antibodies with neutralizing activity. This dataset is unique among viruses in terms of its exceptional size, and thus offers a unique opportunity to separate a polyclonal response into many individual components and then study escape from each one.

For each of 1,603 antibodies from Cao et al., we used the DMS data to estimate an antibody-escape score for each of thousands of globally circulating SARS-CoV-2 sequences from the first 3.5 years of the pandemic. For each antibody, we then computed an escape trajectory quantifying how the average escape score changed in the viral population over time. The trajectories follow several major patterns that differ in terms of their shape and the timing of escape. We validated these patterns using the neutralization data from Cao et al. and Wang et al. [4]. Some trajectories underwent large fluctuations over time as a result of clade-displacement events that reduced the frequency of escape mutations in the viral population. Fitness effects of mutations estimated from sequences in nature suggest that the displacement of escape mutations was often due to clonal interference. The fitness effects also suggest that changes in selective pressure impacted the order in which escape mutations arose.

## Results

### Estimating levels of antibody escape over time

Of the ∼3,000 antibodies characterized by Cao et al., we analyzed a set of 1,603 antibodies that were filtered to only include ones isolated from humans exposed to SARS-CoV-2 (as opposed to SARS-CoV-1) and ones for which the DMS experiments detected strong escape mutations (see *Methods*). The antibodies were isolated from people with a variety of exposure histories, including vaccination with the wildtype (WT)-based vaccine, infection with WT variants, or WT vaccination followed by breakthrough infections with BA.1, BA.2, or BA.5 variants (Table S1), providing a sizable sample of the human polyclonal antibody response during this window in time. As described more below, the antibodies target several epitopes and have a range of abilities to neutralize pseudoviruses bearing the D614G spike variant.

For each antibody, we sought to estimate how levels of antibody escape changed among globally circulating SARS-CoV-2 viruses over time. We did so using the following approach, illustrated for a single example antibody in Figure 1. As input, we used the DMS data from Cao et al. For each antibody, these data measure the effects of thousands of amino-acid mutations spanning the length of the Wuhan-Hu-1 RBD on escape from antibody binding. Specifically, they displayed a mutant library of RBD variants on the surface of yeast cells (where each cell expresses a single RBD variant), incubated the cells with a given antibody, depleted antibody-bound cells from the library using magnetic-activated cell sorting, and then deeply sequenced the library before and after the depletion step to quantify how much each variant escaped antibody binding. From these data, they computed an escape score for each mutation, with scores ranging from 0-1, where 0 indicates that the mutation did not confer escape and 1 indicates that the mutation conferred strong escape. Figure 1A shows the distribution of escape scores for an example antibody. Also as input, we used Nextstrain [35] to obtain a set of 3,942 SARS-CoV-2 genome sequences that were sampled from many countries around the globe and roughly uniformly over time during the first 3.5 years of the pandemic (Figure 1B).

**Figure 1:**
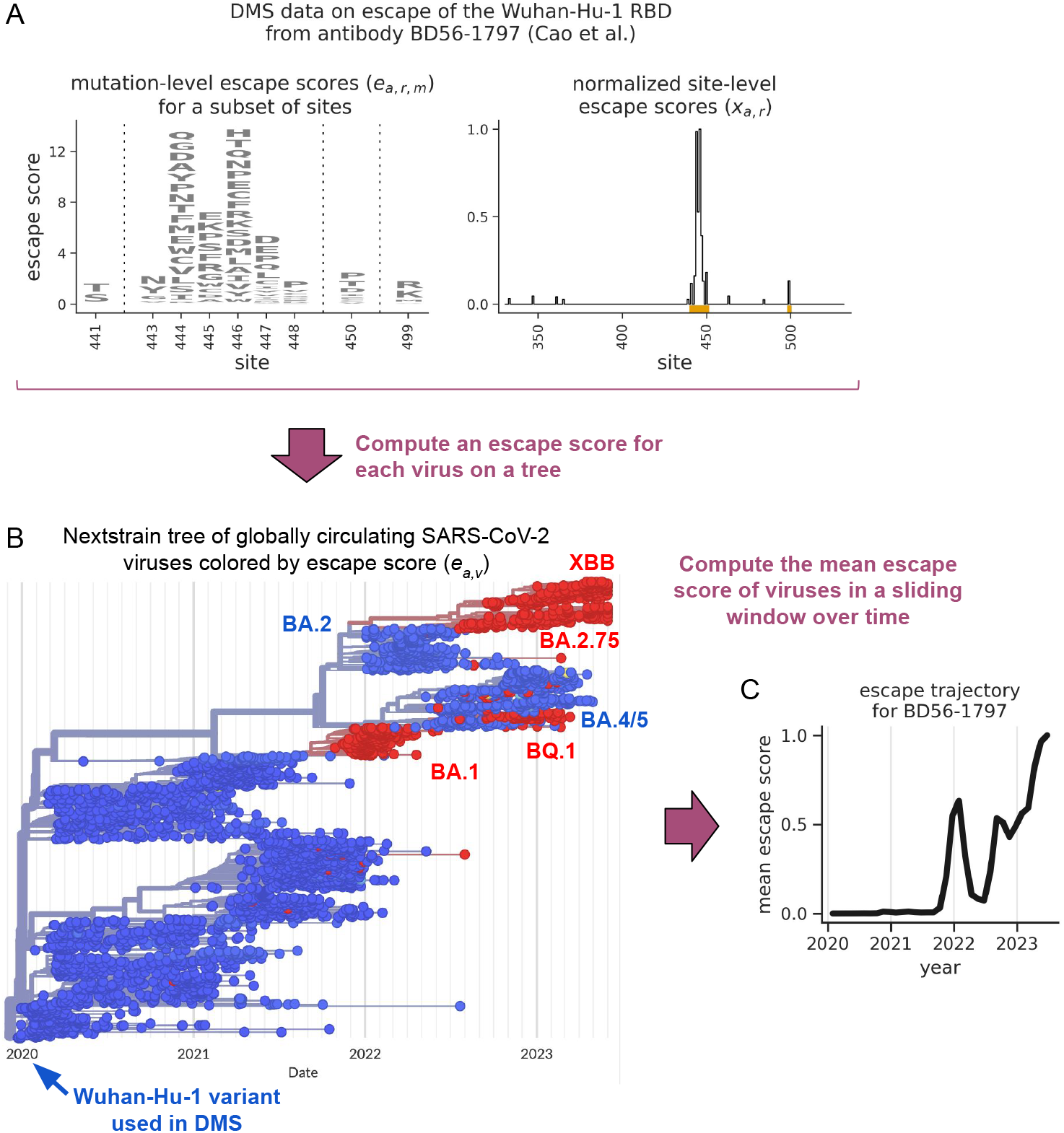
Strategy to estimate levels of antibody escape over time, illustrated for a single example antibody. **(A)** DMS data from Cao et al. quantifying the effects of mutations to the Wuhan-Hu-1 RBD on its ability to escape binding by an example antibody, BD56-1797. The left plot shows amino acid-level escape scores (*e*_*a,r,m*_) for the subset of sites where scores sum to >1. The right plot shows the normalized site-level escape scores (*x*_*a,r*,_), with orange rectangles identifying the sites shown in the left plot. **(B)** A time-resolved Nextstrain phylogenetic tree of all 3,942 globally circulating SARS-CoV-2 viruses under analysis, colored by each virus’s escape score (*e*_*a,v*_) against antibody BD56-1797, as computed from *x*_*a,r*_ values. The tree is colored using a continuous blue-red color scale that ranges from 0-1, with most scores being close to either 0 or 1. **(C)** A 0.2-year sliding-window average of the escape scores shown in panel B. This curve defines SARS-CoV-2’s “escape trajectory” for antibody BD56-1797. We separately compute escape trajectories for each of the 1,603 antibodies under analysis.

For each antibody, we used the DMS data to estimate an escape score for each of the 3,942 viral sequences from above, doing so using a modified version of the strategy from Greaney et al. [36]. Specifically, if *e*_*a,r,m*_ is the escape score of the amino-acid mutation *m* at site *r* of the RBD against antibody *a*, we quantify how much mutating that site tends to escape the antibody as:

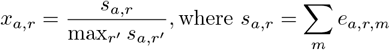

such that that maximum *x*_*a,r*_ value across all sites is always equal to one for a given antibody. Next, we let *e*_*a,v*_ quantify the escape score of virus *v* from the antibody as:

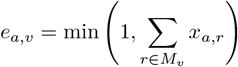

where *M*_*v*_ is the set of sites that are mutated in the virus’s RBD relative to the Wuhan-Hu-1 RBD used in the DMS experiments. We cap *e*_*a,v*_ values at a maximum of one, as values of one already indicate strong escape. Figure 1B shows virus-level escape scores for the example antibody mapped onto a time-resolved tree of the viruses. Computing these scores using *x*_*a,r*_ values is imprecise because it assumes that each amino-acid mutation at site *r* has the same effect. However, as in Greaney et al. [36], we used *x*_*a,r*_ values since they tend to be less noisy than *e*_*a,r,m*_ values. This strategy also assumes, out of necessity, that the effects of all mutations in a viral variant combine additively, i.e., it assumes there is no epistasis between mutations. But, despite these assumptions, we show below that the escape scores correlate with experimentally measured pseudovirus neutralization data.

Finally, for each antibody, we quantified how the average escape score in the viral population changed over time in a 0.2-year sliding window. This sliding-window average, illustrated in Figure 1C for the example antibody, defines the virus’s “escape trajectory” for a given antibody, and is a core part of our subsequent analyses.

Note: the Cao group has also published datasets that characterize antibodies elicited by infections later in the pandemic [17, 37]. The reason why we do not include these datasets in our analysis is that the associated DMS experiments were performed in genetic backgrounds of more evolved RBD variants, rather than the Wuhan-Hu-1 RBD variant. By excluding these additional datasets, we ensure that all viral escape scores are relative to Wuhan-Hu-1.

### Antibody-escape trajectories follow several main patterns

As expected, when we averaged the escape trajectories across all 1,603 antibodies, the resulting average trajectory showed a gradual increase in escape score over time, with the increase starting at about a year into the pandemic when the first variants of concern began to spread (Figure 2A). The average trajectory only reaches a maximum value of ∼0.5, as a subset of antibodies are not predicted to be strongly escaped. This subset almost entirely consists of antibodies with little-to-no neutralizing activity, as described below.

**Figure 2:**
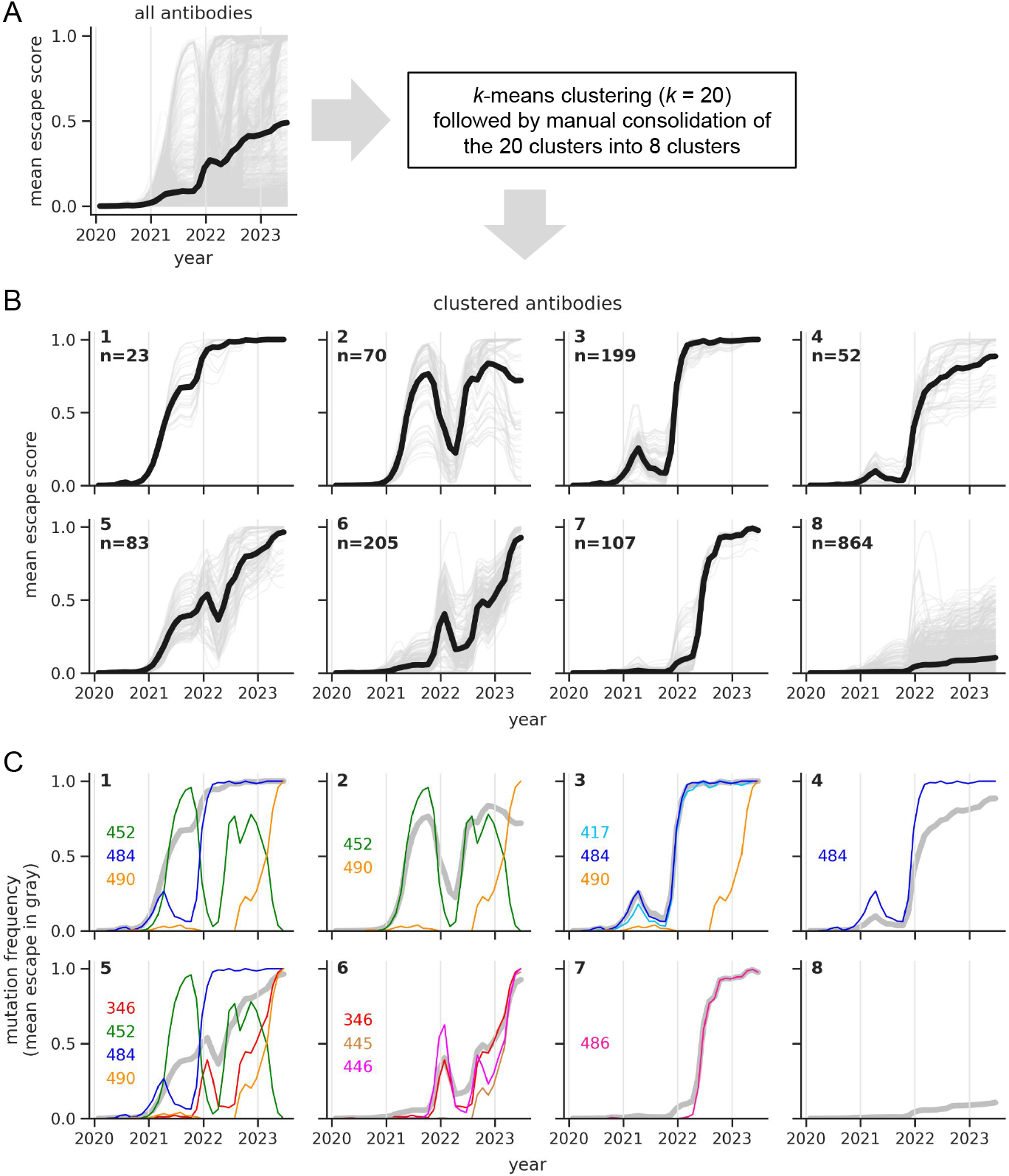
Clusters of antibody escape trajectories and the mutational dynamics underlying escape. **(A)** Average escape trajectory across all antibodies (bold black line) and trajectories for individual antibodies (light gray lines). **(B)** Escape trajectories partitioned into clusters, with the bold black line showing the cluster average and the light gray lines showing trajectories for individual antibodies. The top left corner of each plot reports the cluster label (1-8; clusters are roughly ordered according to when the virus escaped them) and the number of antibodies within each cluster. **(C)** For each cluster, the frequencies of mutations at sites that are key drivers of the escape trajectories (narrow colorful lines), along with the average escape trajectory for a given cluster, as in panel B (bold gray line). Mutational frequencies are computed using the same 0.2-year sliding windows as the trajectories.

In contrast, when we examined escape trajectories of individual antibodies, we saw a variety of trends. To identify these trends, we first clustered the trajectories into 20 clusters using *k* -means clustering, which revealed fine-grained trends (Fig. S1). We then manually consolidated these 20 clusters into 8 clusters that highlight interesting biological patterns in the data (Figure 2B). Average escape trajectories differ between clusters in a few key ways. First, they differ based on the timing of escape, with different clusters showing sequential waves of escape that span multiple years. Second, they differ in their shape. Some trajectories steadily increase over time before plateauing at a high escape score (e.g., clusters 1 and 7), while others show large fluctuations, where the escape score starts to increase, then substantially decreases before increasing to a high escape score (e.g., clusters 2, 3, 6). The trajectories in cluster 8 show no increase in escape score or plateau at values well below one. Individual trajectories are somewhat continuous between certain pairs of clusters (e.g., clusters 3 and 4) due to continuity in antibody DMS profiles [3].

In the next three sections, we examine the escape mutations that gave rise to these patterns, we examine the epitopes and neutralization activity of antibodies from each cluster, and we validate cluster-averaged escape patterns using experimental pseudovirus neutralization data.

### Some escape trajectories arose through complex mutational dynamics

For each cluster, we sought to identify key sites that drove escape. We did so by searching for sites that had an intermediate-to-high escape score (*x*_*a,r*_ > 0.4) for at least a quarter of the antibodies in a given cluster, and where mutations reached a high frequency (>0.9) in nature. Figure 2C lists the sites that met these criteria for each cluster, and shows how changes in mutational frequencies at these sites drove each cluster’s average escape trajectory.

For some clusters, the average escape trajectory closely tracks with the frequency of mutations at a specific site. For instance, the average escape trajectory from cluster 7 closely tracks with the frequency of mutations at site 486, while the average escape trajectory from cluster 3 closely tracks with the frequency of mutations at either 417 or 484. The reason that the cluster 3 trajectory tracks with two different sites is that antibodies from this cluster tend to be strongly escaped by mutations at either 417 or 484 (Fig. S2; compare cluster 3 epitopes A and C), and escape mutations at these sites are genetically linked, such that both groups of antibodies have similar escape trajectories.

In contrast, for other clusters, the average escape trajectory does not just track with the frequency of mutations at a single site, but is instead shaped by a combination of escape mutations at multiple sites in the same antibody epitope. Often, these escape mutations undergo large fluctuations over time, with one strong escape mutation declining in frequency while another rises, such that escape is maintained over time by different mutations (see clusters 1, 2, and 5). Some sites are key drivers of escape for multiple clusters. This results from antibodies from different clusters having overlapping, but distinct epitopes (Fig. S2). For instance, clusters 1 and 2 share two key sites where mutations drive escape (sites 452 and 490), but differ in a single key site (484). Similar patterns are apparent for clusters 1 and 3 and clusters 5 and 6. These subtle differences in antibody epitopes give rise to large differences in escape trajectories.

### Antibody epitopes and neutralization potencies

The antibodies in our analysis target several epitopes, defined by Cao et al. based on clustering of their DMS profiles, and have a range of abilities to neutralize pseudoviruses bearing the original Wuhan-Hu-1 D614G spike variant (Figure 3A). As might be expected, antibodies that target the same epitope often fall in the same escape cluster (Figure 3B), but not always, such that clustering trajectories by epitope conflates certain escape patterns (Fig. S3). Sometimes, antibodies targeting *different* epitopes fall in the same cluster, either due to escape mutations in different epitopes being genetically linked or, in the case of cluster 8, due to little-to-no predicted escape.

**Figure 3:**
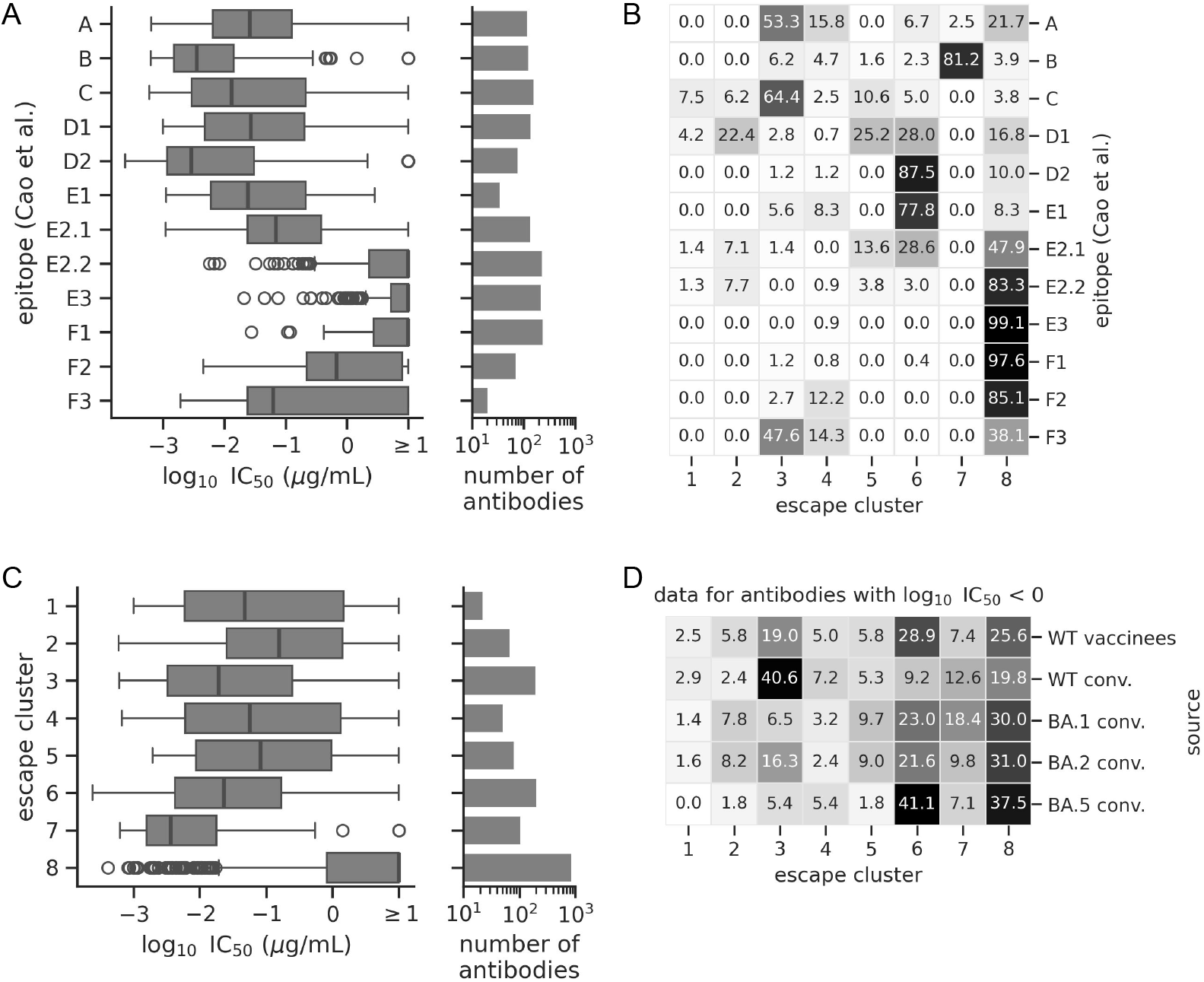
Antibody epitopes and neutralization potencies. **(A)** Data for the 1,603 antibodies grouped by epitope (defined by Cao et al.), with the left plot showing the distribution of log_10_ IC_50_ values quantifying the ability of an antibody to neutralize pseudoviruses bearing the D614G spike, and the right plot showing the number of antibodies within each epitope. **(B)** A heat map showing the percent of antibodies in a given epitope that are in a given escape cluster (Figure 2B), such that each row sums to 100. **(C)** Same as panel A, but showing data for antibodies clustered by escape trajectory. **(D)** A heat map showing the percent of antibodies from a given source that are in a given escape cluster, such that each row sums to 100 (conv. = convalescent). The plot only shows data for antibodies with appreciable neutralizing activity against the D614G variant (log_10_ IC_50_ < 0 *µ*g/mL).

The antibodies from clusters 1-7, which are predicted to be strongly escaped, often neutralize the D614G variant, while those from cluster 8, which are not predicted to be strongly escaped, tend to have little-to-no neutralizing activity (Figure 3C). This pattern is consistent with the expectation that neutralizing antibodies exerted the greatest selection on the virus.

However, the antibodies escaped first were not always the most potent. Antibodies from clusters 3, 6, and 7 stand out as having especially high neutralization potency and abundance (Figure 3C). Cluster 3 antibodies are predicted to be among the first that were completely escaped, as defined by trajectories reaching values close to one (Figure 2B). However, cluster 6 and 7 antibodies were among the last to be completely escaped, raising the question of why these antibodies were not escaped sooner.

One hypothesis is that antibody abundance changed over time. Grouping antibodies by host source shows that neutralizing antibodies from WT exposures, which occurred earlier in the pandemic, are particularly enriched in cluster 3, while those from the BA.5 infections, which occurred later, are particularly enriched in cluster 6 (Figure 3D). Thus, shifts in immunity may partially explain the order of escape. However, aside from clusters 3 and 6, the distribution of neutralizing antibodies between clusters is largely similar between host sources (Figure 3D), likely due to imprinting effects described by Cao et al. [3]. Thus, factors other than antibody potency and abundance may have contributed to the order of escape. We investigate this possibility more below.

### Validation of average escape patterns from each cluster using pseudovirus neutralization data

We next sought to validate cluster-averaged patterns using pre-existing neutralization data from Cao et al. and Wang et al. [4]. For each antibody from Cao et al., the authors measured the ability of the antibody to neutralize pseudoviruses bearing one of several spike variants, including the D614G variant and several Omicron variants of concern. For a small subset of these antibodies, Wang et al. used the same approach to measure the ability of each antibody to neutralize the D614G variant, several pre-Omicron variants of concern, and BA.1. Together, these spike variants are widely distributed across the tree of viruses in our analysis, are founder sequences of large clades, and capture large changes in predicted escape scores, making them well-suited for validating predicted trends.

The above data were collected using large-scale neutralization assays, and as such, have a non-trivial level of noise. We quantified this noise by examining the correlation of log_10_ IC_50_ values between Cao et al. and Wang et al. for antibody-variant combinations measured in both studies. These values were correlated (Pearson R = 0.87), but often differ by one or more log_10_ units (Figure 4A).

**Figure 4:**
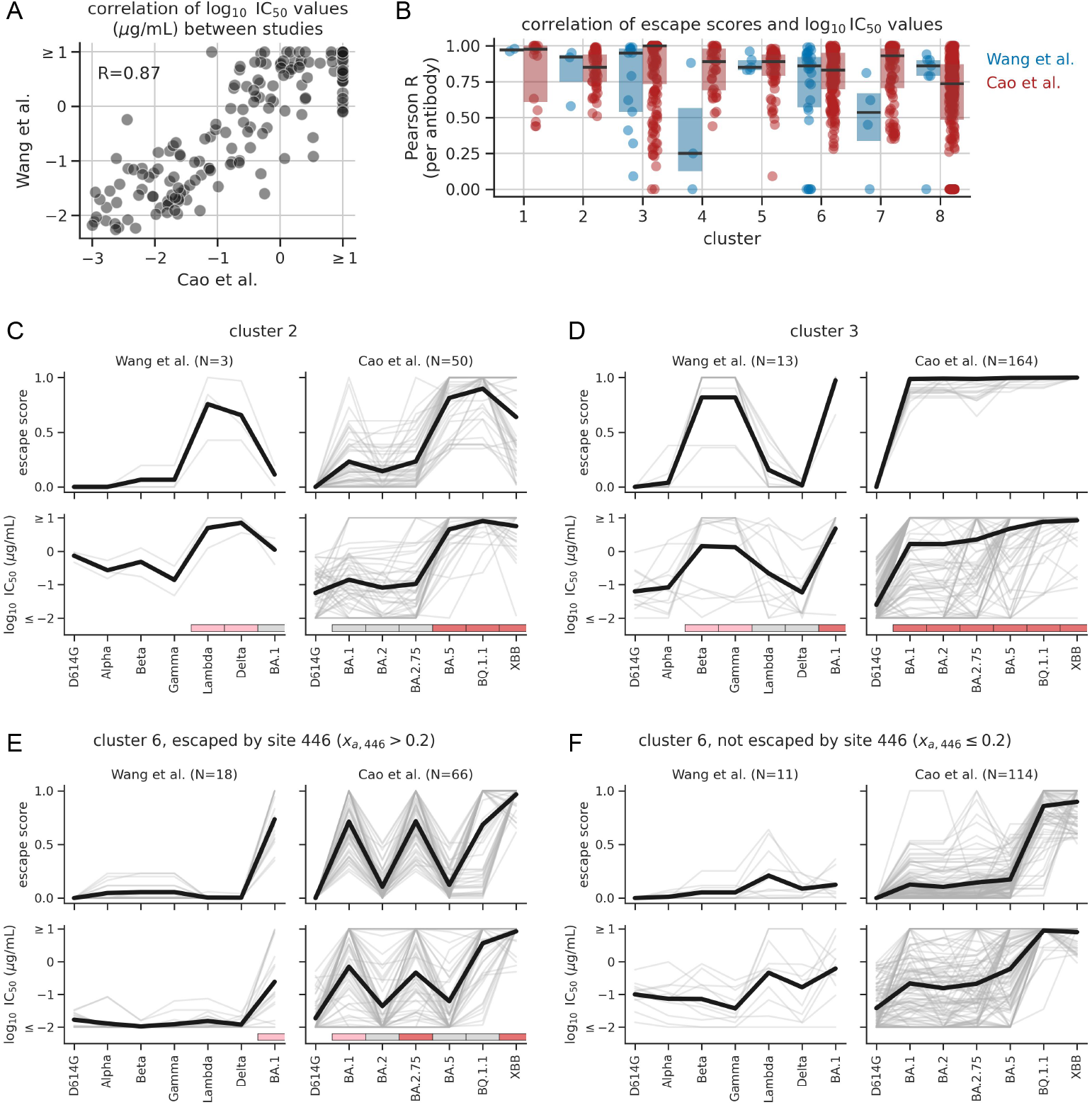
Predicted escape scores correlate with pseudovirus neutralization data. **(A)** Correlation of log_10_ IC_50_ values between studies. **(B)** Distributions of perantibody Pearson correlation coefficients quantifying the agreement between predicted escape scores and log_10_ IC_50_ values for spike variants from a given study. Each dot corresponds to a single antibody and antibodies are grouped by cluster. Shaded boxes show interquartile ranges, and horizontal black lines show medians. We applied a floor of zero to each distribution. **(C)-(F)** Comparison of escape scores and log_10_ IC_50_ values for antibodies from a given cluster. In each panel, the left column of plots shows antibodies with neutralization data from Wang et al., the right column shows antibodies with neutralization data from Cao et al., and N indicates the number of antibodies with data. The x-axis shows the spike variants assayed in each study, with variants ordered roughly chronologically by when they emerged in nature. Bold black lines show average values, while thin gray lines show values for individual antibodies. The log_10_ IC_50_ values have an upper limit of detection of 1, and we floored the values at a lower limit of −2. Panels E and F show data for two groups of antibodies in cluster 6: ones escaped by mutations at site 446 (*x*_*a*,446_ > 0.2) and ones that are not (*x*_*a*,446_ ≤ 0.2). In panels C-E, the pink, gray, and red rectangles below the neutralization data correspond to the cluster-specific clade groupings from Figure 5. Fig. S4 shows similar plots for the remaining clusters.

We found that the neutralization data are consistent with many of the cluster-averaged patterns of escape. We specifically focused on antibodies with appreciable neutralizing activity against the D614G variant (log_10_ IC_50_ < 0 *µ*g/mL), since the neutralization data are only suitable for validating escape from neutralizing antibodies. Figure 4C-F show data for the three clusters with the largest fluctuations in average escape trajectories (clusters 2, 3, and 6). The predicted escape scores show striking increases and decreases between adjacent spike variants when the variants are ordered roughly by when they emerged. When averaging over all antibodies in a cluster, the changes in escape score are mirrored by changes in log_10_ IC_50_ values (see the bold black lines). At the same time, for a subset of individual antibodies, the changes are clearly not mirrored, either due to inaccurate predictions, experimental noise, or experimental limits of detection.

The antibodies from cluster 6 show two distinct neutralization profiles depending on whether the antibody is strongly escaped by mutations at site 446 or mutations at another site, most commonly site 346 (Figure 4E and F, respectively). The key difference is that mutations at site 446 are present in the BA.1 and BA.2.75 variants, while mutations at site 346 are not. These two groups of antibodies still have similar predicted escape trajectories because mutations at site 346 reach high frequencies within the BA.1 and BA.2.75 clades, even though they are not present in the clade-founder sequences used in the neutralization assays (we describe this pattern in more detail below).

To quantify levels of agreement, for each antibody, we computed the Pearson correlation coefficient between escape scores and log_10_ IC_50_ values among all spike variants from a given study (Figure 4B). For each cluster of antibodies, the resulting median correlation coefficient was often between 0.75-1.0, with only a small subset of individual antibodies reaching low correlation coefficients. The median correlation coefficient for the Wang et al. study was low for cluster 4 and 7, though these medians are only derived from a total of 3 and 4 antibodies, respectively, and the average trajectories of these clusters, which are largely shaped by Omicron variants, are supported by the Cao et al. data (Fig. S4). Unfortunately, most clusters only had a few antibodies with validation data from Wang et al., limiting our ability to robustly validate all trends using these data. But, overall, the data are still consistent with many of the cluster-averaged escape patterns.

One reason that the above correlation coefficients are high is that both escape score and neutralization resistance tend to generally increase with sequence divergence, regardless of cluster. To quantify this effect, we recomputed the correlation coefficients using the number of RBD mutations in a given variant in place of its escape score. For clusters 1-3 and 5-7, the resulting coefficients are often substantially lower than before for at least one study (Fig. S5), indicating that escape scores capture more than just sequence divergence.

The validation data are difficult to interpret for cluster 8 antibodies. For this cluster, we do not find evidence that escape scores capture more than sequence divergence (Fig. S5). Further, since most escape scores from this cluster only slightly increase over time, a concern is that the increase is merely due to the summed effects of noise from the DMS experiment, such that the correlation with the neutralization data is artificial. Despite the modest increases in predicted escape score, there is a large increase in average log_10_ IC_50_ values between the D614G variant and the BQ.1.1 and XBB variants (Fig. S4), indicating that the DMS experiments may not have identified relevant escape mutations, possibly due to experimental noise, or possibly because the relevant mutations do not confer escape in the Wuhan-Hu-1 background. These data do not shed light on whether the virus also escaped *non*-neutralizing antibodies from cluster 8, which constitute the majority of antibodies from this cluster.

### Large dips in escape trajectories track with clade-displacement events

Clusters 2, 3, and 6 show large dips in escape trajectories. As described above, these dips happen as a result of escape mutations decreasing in frequency in the viral population. These decreases are due to clade-displacement events. Figure 5A-C illustrates this phenomenon in context of a single example antibody from each cluster. In each case, the antibody escape trajectory begins to increase as an escape mutation rises in frequency in the viral population. The mutation is initially found in the group of clades colored in pink in the figure. Next, the gray group of clades, which lack the mutation, displace the pink group. This causes a dramatic decrease in the overall frequency of the escape mutation, and thus the escape trajectory. Finally, the red group of clades, which have escape mutations, displace the gray group, causing the frequency of these mutations and the trajectory to increase again. (Observe the concordance between the “mean escape” trajectory and the union of the pink and red clade frequency trajectories.) We see similar patterns with other escape mutations associated with fluctuating trajectories (Fig. S6A-B).

**Figure 5:**
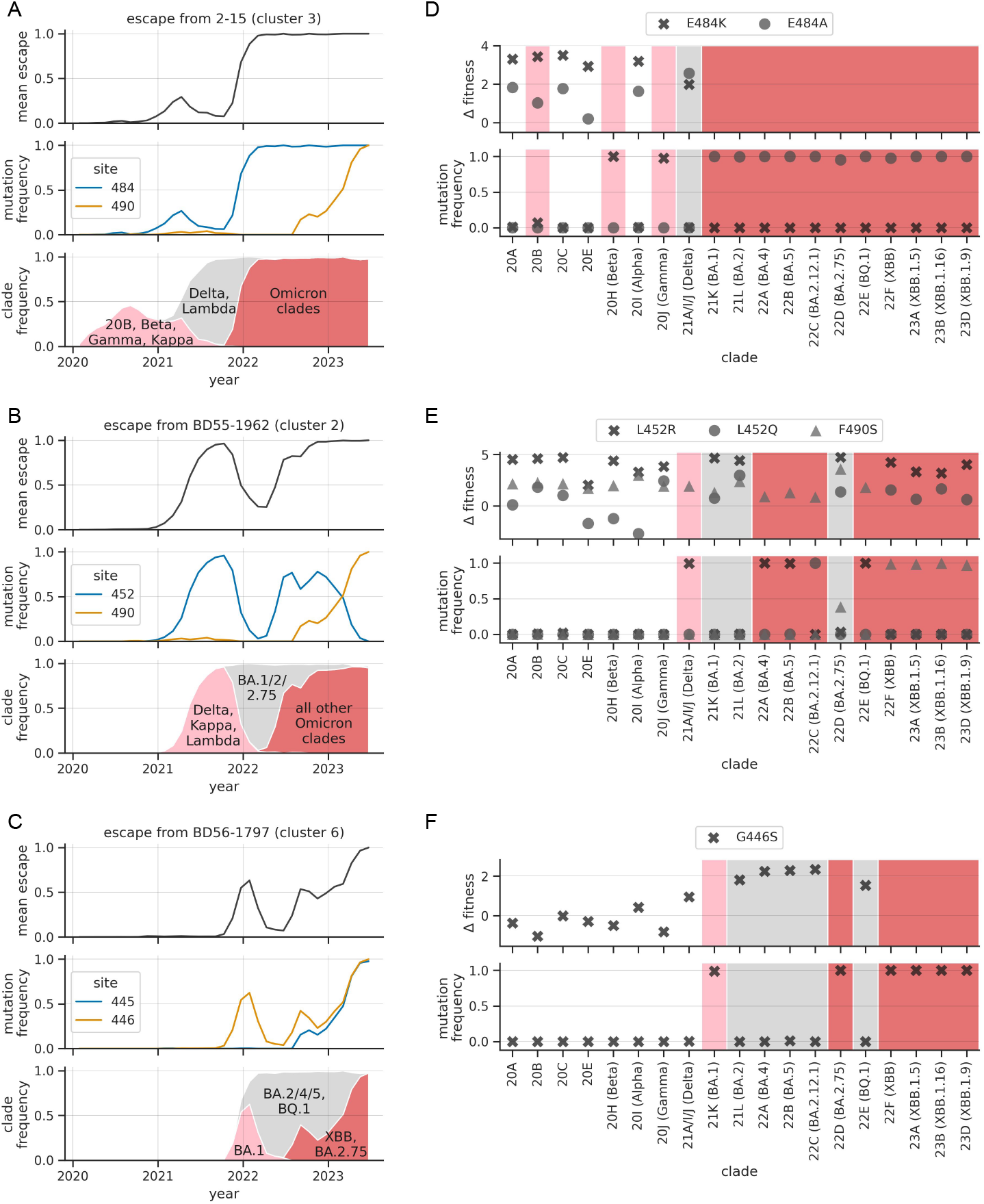
Clade-displacement events cause large dips in trajectories. **(A)-** Each panel shows data for an example antibody from a given cluster. For each antibody, rows show the escape trajectory for that antibody, the frequency of mutations at key sites of escape (i.e., sites where *x*_*a,r*_ > 0.4 and where mutations reach a frequency of >0.9), and the frequencies of groups of clades involved in displacement events that shape a given trajectory. Clades in the pink and red groups have escape mutations at appreciable frequencies, while clades in the gray group do not. Trajectories dip when gray clades displace pink clades and rise when red clades displace gray clades. We use Nextstrain clade definitions and compute frequencies using the same 0.2-year sliding windows used to compute the trajectories. **(D)-(F)** Each panel shows data for specific amino-acid mutations that drove escape trajectories in panels A-C (compare panels in the same row). In each panel, the bottom plot shows mutation frequencies in each of the indicated Nextstrain clades (with parenthetical names giving the variant of concern that corresponds to a clade’s founder sequence, when applicable). For simplicity, we only show data for clades that reached >7% frequency. Clades are colored according to how they are grouped in panels A-C. The top plot shows estimates of mutation fitness effects in each clade (effects are not estimated for clades that already have a mutation at a given site).

The trajectories closely track with clade-displacement events because the frequency of escape mutations within individual clades is often either close to zero or one (see the bottom plot of each panel of Figure 5D-F; the upper plot will be described in the next section). An exception to this trend is site 346, where escape mutations have intermediate frequencies in multiple clades (Fig. S6E).

In the examples from Figure 5, the clades that caused the trajectory of one antibody to decrease often caused the trajectory of another to increase. The spread of Delta and Lambda caused the cluster-3 antibody trajectory to decrease, but caused the cluster-2 antibody trajectory to increase. The spread of BA.1 caused the cluster-2 antibody trajectory to decrease, but caused the cluster-3 and cluster-6 antibody trajectories to increase. Such patterns suggest that clade-displacement events resulted in large tradeoffs in resistance to different clusters of antibodies, despite the fact that the virus’s overall antibody resistance steadily increased over time.

### Evidence that clonal interference caused the displacement of escape mutations

We next sought to investigate why clade-displacement events were effective at driving down the frequencies of escape mutations in the viral population. SARS-CoV-2 can undergo genetic recombination [38], which provides a mechanism to consolidate beneficial mutations from different genetic backgrounds [39]. In each example from Figure 5A-C and Fig. S6A-B, recombination would have offered a mechanism to introduce the escape mutation from the pink clades into the genetic background of the gray clades, helping to prevent the escape mutation from being displaced. According to our additive model, such recombination events would have been beneficial, as they would have increased escape scores of viruses in the grey clades. It is possible that such recombination events did occur at a low level. But, ultimately, none of the displaced mutations reached appreciable frequencies in the gray clades (Figure 5D-F; S6D-E).

There are multiple possible explanations for this pattern. One explanation is clonal interference. For organisms that do not undergo genetic recombination, clonal lineages with different beneficial mutations can compete with one another, delaying the rate at which beneficial mutations are fixed in the population [40, 41, 42, 43]. Although SARS-CoV-2 does recombine, it is possible that recombination has not always been efficient enough to prevent competition between clades from displacing beneficial mutations. Another possible explanation is differences in selective pressures between clades. The escape mutations in the above examples may have had beneficial fitness effects in the pink clades, but neutral or deleterious effects in the gray clades. This could happen due to epistasis (the mutation is incompatible in the genetic backgrounds of the gray clades) or due to changes in external selective pressures (e.g., if the strength of selection imposed by a given cluster of antibodies varied over time), such that even if the mutation was recombined into a gray clade, it would not have conferred a selective advantage.

To investigate these possibilities, we used a previously published computational pipeline to estimate fitness effects of mutations from natural sequences [44, 45]. As input, we used a phylogenetic tree of ∼16 million SARS-CoV-2 genome sequences. The pipeline estimates a mutation’s effect based on the number of times the mutation was observed to arise *de novo* on the branches of the tree versus the number of times it was expected to have arisen under neutral evolution. Specifically, it estimates the effect as the natural log of the ratio of observed counts to expected counts, such that mutations that arose more than expected have positive (i.e., beneficial) fitness effects, and vice versa for mutations that arose less than expected. It performs this estimation using a Bayesian approach that integrates over uncertainty in the observed and expected counts. A mutation’s effect is defined as the mean of the posterior probability distribution over all possible effects. In addition, the standard deviation of this distribution can be used to quantify the level of uncertainty associated with a given effect. Uncertainty is modeled to arise from statistical sampling noise in the counts, as well as unaccounted variability in neutral mutation rates between sites. We used this approach to estimate mutational effects at the level of individual clades in our analysis.

Under the clonal-interference hypothesis, we would expect escape mutations from the pink clades to have positive fitness effects in the gray clades that displace them. Indeed, we see this pattern for four out of the six displaced escape mutations. For example, as shown in Figure 5F, the G446S escape mutation, which is fixed in the pink BA.1 clade (see the bottom plot), has large positive fitness effects in the corresponding gray clades (see the top plot). These large positive effects are much larger than associated levels of uncertainty (each effect is >2-fold larger than the standard deviation of the corresponding posterior distribution; Fig. S7C), providing confidence that the effects are indeed positive. This suggests that the G446S mutation was beneficial in the gray clades, but that recombination of this mutation into sequences from the gray clades was not efficient enough to prevent its displacement. We see similar evidence of clonal interference for E484K (Figure 5D; see the large positive effect of this mutation in the Delta clade), L452R (Figure 5E; see the large positive effect of this mutation in the BA.1, BA.2, and BA.2.75 clades), and R346K (Fig. S6E; see the large positive effect of this mutation in the BA.2 clade). As above, these large positive effects are much larger than associated levels of uncertainty (Fig. S7). For R346K, we also find support for changing selective pressures between clades: the mutation is present at high frequency in the pink BA.1 clade and has a large positive fitness effect in the gray BA.2 clade (suggesting clonal interference), but has more modest positive effects in subsequent gray clades (suggesting changing selective pressures).

For the four mutations that show signs of clonal interference, their fitness effects in the gray clades range from ∼2-5 units, meaning that the substitution rate of these mutations was ∼7-150 times higher than expected under neutrality. Despite these high substitution rates, most mutations did not reach frequencies appreciably greater than zero within the gray clades. This apparent discordance is because substitution rates were estimated from millions of viral sequences, and suggests that although mutations may arise *de novo* at a high rate, it takes time for them to reach appreciable frequencies in the very large population.

The remaining two displaced escape mutations (K417N and K417T) show little or no evidence of clonal interference. Each mutation has a roughly neutral fitness effect in the gray Delta clade that displaced it (Fig. S6D; Fig. S7D).

### Evidence that changing selective pressures impact the order in which escape mutations arose

The key sites of escape from Figure 2C can be divided into two broad groups based on whether or not escape mutations at a site reached appreciable frequencies before the emergence of Omicron in late 2021. They did at sites 417, 452, and 484, and did not at sites 346, 446, 486, and 490. Why did some escape mutations not appear before the emergence of Omicron?

The fitness data from the previous section indicate that the effects of some mutations shifted to become dramatically more beneficial upon the emergence of Omicron. We see this pattern for multiple key escape mutations at the second group of sites: R346T, G446S, and F486V (Figure 5F; Fig. S7E-F). In pre-Omicron clades (20A-21J), each of these mutations has a roughly neutral effect, while in Omicron clades (21K-23D), each has a large positive effect. These positive shifts in fitness effects are greater than associated levels of uncertainty (Fig. S7). Thus, these shifts could help explain why these three antibody-escape mutations only arose after the emergence of Omicron.

The above shifts could be due to multiple factors, including shifts in human immunity over time or shifts in the effects of mutations on spike’s function or ability to escape antibodies, due to epistasis with pre-existing mutations in Omicron clades. Indeed, past work has identified multiple examples of mutations with enhanced antibody-escape effects in Omicron backgrounds [46, 47], including mutations at sites 346, 446, and 486. It is possible that such enhancement could explain the increased fitness of the above mutations in Omicron clades. Past work has also identified examples of escape mutations, such as G446S, that are deleterious to RBD-ACE2 binding in context of the Wuhan-Hu-1 RBD, but are buffered by other affinity-enhancing mutations in Omicron BA.1 [48, 49]. When we used data from Starr et al. [48, 46] to comprehensively examine the ACE2-binding effects of each of the key antibody-escape mutations from our analysis, we found that mutations at 417, 446, and 486 tended to have deleterious effects, while the mutations at other sites tended to have roughly neutral effects (Fig. S8). If variants from Omicron clades are generally better at buffering deleterious effects, then that could explain why both G446S and F486V have increased fitness effects in these clades. Further, while G446S is deleterious to ACE2 binding in pre-Omicron variants, its effect is much less deleterious in the BA.2 variant (Fig. S8), indicating a reduced need for buffering in this Omicron variant and potentially in others.

In all, the above patterns suggests that there are multiple factors that can determine how quickly antibody-escape mutations appear and then fix in the viral population, including both clonal interference and changes in underlying selective pressures over time.

## Discussion

Leveraging the DMS data from Cao et al., we predicted antibody-escape scores for thousands of globally circulating viral variants, doing so for each of 1,603 antibodies. For each antibody, we then quantified how the average escape score changes in the viral population over time. We distilled the resulting trajectories into eight major clusters, which show sequential waves of escape.

Although some trajectories steadily increase in escape score over time, others show striking fluctuations. We validated cluster-averaged trajectories by showing that predicted escape scores correlate with pseudovirus neutralization data for several variants of concern (using neutralization data from Cao et al. and Wang et al.). We showed that fluctuations in trajectories track with clade-displacement events that displaced key escape mutations from the population. Phylogenetically estimated fitness effects suggest that the mutations were often displaced due to clonal interference. The fitness estimates also indicate that the order in which escape mutations arose is shaped by changing selective pressures. Our work builds on past studies of SARS-CoV-2 antibody escape in a few ways. First, we examined escape from a much larger number of antibodies than most studies. This allowed us to catalogue many different patterns of escape in a single study, and to provide a more comprehensive picture of the fraction of antibodies that follow each pattern. For instance, past studies have also identified antibodies for which levels of resistance fluctuated between variants of concern over time, including antibodies with neutralization profiles that are similar to those of clusters 2, 3, and 6 from Figure 4. R1-32 and FC08 have profiles that are similar to cluster 2 and are also escaped by mutations at site 452 [16, 50]; LY-CoV016 (etesevimab) and REGN10933 (casirivimab) have profiles that are similar to cluster 3 and are also escaped by mutations at sites 417 and 484 [33, 51]; REGN10987 (imdevimab), C110, and 2H04 have profiles that are similar to cluster 6 and are also escaped by mutations at site 446 [52]. Building on these studies, our work indicates that a substantial fraction of the hundreds of neutralizing antibodies from our analysis have fluctuating escape trajectories. Thus, although our estimates indicate that the virus’s average antibody resistance tended to steadily increase over time, they also indicate that the underlying escape patterns were complex, with large tradeoffs in resistance between different groups of antibodies.

Second, we established a framework to use DMS data to estimate how levels of antibody resistance changed over time in the global SARS-CoV-2 population. Because of the low-throughput nature of classical neutralization assays, past studies have typically only measured escape for several viral variants of concern. While experimental measurements would tend to be more accurate, the DMS-based approach offers quantitative estimates of population-level escape over time. These estimates suggested a pattern that was not evident in the neutralization data from Cao et al. and Wang et al.: antibodies in cluster 6 show one of two distinct neutralization profiles, but are predicted to have similar escape trajectories due to the emergence of mutations at site 346 within several clades. Further, with the DMS-based estimates, it is straightforward to identify which mutations were predicted to drive escape trajectories, allowing us to hypothesize how trajectories arise from the dynamics of specific mutations and clade-displacement events.

Third, we uncover evidence that clonal interference delays the fixation of antibody-escape mutations in SARS-CoV-2’s evolution. This phenomenon has been observed for viruses that do not genetically recombine [53, 54, 55]. However, it has been unclear whether it impacts SARS-CoV-2. Recombination has been detected among natural SARS-CoV-2 isolates [38], including an event that gave rise to the XBB variant of concern [39], which spread globally. Thus, SARS-CoV-2 recombination can produce highly fit variants. Building on this work, our findings indicate that although SARS-CoV-2 recombination is possible, it has not always been efficient enough to prevent beneficial escape mutations from being displaced, leading to clonal interference. In each example of this pattern that we characterized, the displaced mutation (or another at the same site) eventually rebounded in frequency after about six months to a year, suggesting that although clonal interference may have delayed SARS-CoV-2’s antigenic evolution, the delay was not long.

Fourth, our work emphasizes that it can take a substantial amount of time for positively selected mutations to reach appreciable frequencies in the global population. In examining clade-specific mutational fitness effects, we identified several instances where the *de novo* substitution rate of an escape mutation within a clade was ∼7-150 times higher than expected under neutral evolution, indicative of positive selection. But, in most such instances, the overall frequency of the mutation in the clade was still close to zero. Thus, high levels of positive selection do not necessarily lead to large increases in mutation frequency during the lifetime of a clade. This could help explain why recombination does not always prevent clonal interference: even if recombination produces fit variants at a low basal rate, those variants are not guaranteed to immediately become widespread.

Fifth, our work adds support to the hypothesis that changing selective pressures impacted the order in which escape mutations arose. Previous studies have provided compelling experimental evidence that compensatory mutations in the BA.1 genetic background help buffer functionally deleterious effects of other BA.1 mutations that confer antibody escape [48, 49]. Studies have also shown that some mutational effects on antibody escape are potentiated in Omicron backgrounds [46, 47]. Here, we add to this work by showing that fitness effects estimated from natural sequences show dramatic increases in Omicron clades compared to pre-Omicron clades for three key escape mutations from our analysis. And we show how these shifts can help rationalize the order in which key escape mutations from our analysis arose in nature.

Our study has a few limitations. First, the DMS data measure the effects of mutations on RBD escape from antibody binding in a yeast-display system, not effects on escape from antibody neutralization in a viral context. Second, the additive model that we use to predict antibody-escape scores makes several assumptions. It assumes no epistasis between mutations. Though, as discussed above, previous studies have identified examples of strong epistasis in RBD’s evolution. It also assumes that normalized escape scores are directly comparable between antibodies, that all mutations at a site have the same effect, and that variant escape scores have an upper limit of one. The above assumptions, together with noise from the DMS experiments, limit the accuracy of estimated escape scores. Indeed, the validation data suggest that the estimates are incorrect for a subset of antibodies. It is encouraging that the validation data are consistent with cluster-averaged trends, which form the core of our analysis. And it is also notable that similar DMS-based additive approaches have been successful at predicting levels of antibody escape for variants of viral entry proteins from Influenza virus and Lassa virus [56, 57]. However, a third limitation of our study is that some clusters only had a few antibodies with validation data from the Wang et al. study, which limits our ability to robustly validate all trends without additional experiments. Fourth, the mutational fitness effects estimated from naturally occurring sequences are difficult to experimentally validate, and the trends indicating clonal interference and changing selective pressures are mainly supported by four and three mutations, respectively. We will be curious to see if similar trends are apparent in the on-going evolution of SARS-CoV-2.

Overall, our work helps dissect how SARS-CoV-2 evaded the individual components of a polyclonal antibody response in nature. In the future, our analysis framework could be applied to other contexts. DMS is increasingly being used to identify mutations to viral entry proteins that confer antibody escape, both for SARS-CoV-2 [17, 58, 59, 60] and for other viruses [56, 57, 61, 62, 63, 64, 65]. Further, many viruses have pre-existing Nextstrain [35] pipelines for creating curated multiple-sequence alignments from real-time surveillance data (https://nextstrain.org/). Our framework, and others like it, offer an exciting way to combine these sources of data to interpret patterns of viral evolution in nature.

## Methods

### Data and code availability

See https://github.com/matsengrp/ncov-ab-escape for input files, code, and key results files from our analysis. Key results files include:

- a file with metadata on the viral sequences from our analysis (https://github.com/matsengrp/ncov-ab-escape/tree/main/results/viral_metadata.csv).
- a file with site-level escape scores (https://github.com/matsengrp/ncov-ab-escape/tree/main/results/processed_input_data/escape.csv).
- a directory with files with fitness effects estimated from natural sequences (https://github.com/matsengrp/ncov-ab-escape/tree/main/results/aamut_fitness/).
- a file with each antibody’s escape trajectory and escape cluster (https://github.com/matsengrp/ncov-ab-escape/tree/main/results/trajectories.csv).

The README.md file in the repository’s root directory describes the repository’s contents in more detail.

### Input DMS data

We got the input DMS data from Cao et al. [3]. Their repository https://github.com/jianfcpku/convergent_RBD_evolution includes the DMS data for each antibody, as well as other metadata associated with each antibody, including the antibody’s source and its neutralization data against variants of concern. For practical reasons, the set of publicly available escape scores, which we use in this study, were filtered to remove data at sites where all escape scores were close to zero for a given antibody. We assign these filtered-out mutations escape scores of zero for that antibody.

Of the 3,333 antibodies with data, we curated a set of 1,603 using the following filters: i) the antibody was strongly escaped in the DMS experiments, defined as the *e*_*a,r,m*_ values for that antibody summing to >2 for at least one site, ii) the antibody was isolated from a human exposed to SARS-CoV-2 (rather than SARS-CoV-1), and iii) the antibody had pseudovirus neutralization data for all variants from the Cao et al. study from Figure 4. There were 2,326 antibodies that passed the first filter, 1,616 antibodies that passed the first two filters, and 1,603 antibodies that passed all three filters.

### Sampling globally circulating viruses

We sampled 3,942 globally circulating viruses from the first 3.5 years of the pandemic using a modified version of the ncov Nextstrain [35] workflow (https://github.com/nextstrain/ncov). See https://github.com/matsengrp/ncov-ab-escape for the specific code and input files that we used to run this workflow, as well as key output files, including a file that lists for each viral sequence: the set of amino-acid mutations in that sequence’s spike protein relative to the Wuhan-Hu-1 reference strain, the year the sequence was sampled, and the Nextstrain clade associated with the sequence. The repository also includes the phylogenetic tree of these sequences from Figure 1B.

### Computing viral escape scores

We computed viral escape scores as described in the main text. We used the multiple-sequence alignment of spike sequences from above to define the set of mutations in each sequence relative to the Wuhan-Hu-1 reference sequence (called “Wuhan-Hu-1/2019” in the alignment). We only considered mutations in the RBD, since the DMS data were specific to this domain. See https://github.com/matsengrp/ncov-ab-escape for the specific code we used to compute viral escape scores. We adapted this code from the escape calculator (https://github.com/jbloomlab/SARS2-RBD-escape-calc/) initially described in Greaney et al. [36].

### Clustering of escape trajectories

We used *k* -means clustering to cluster trajectories into 20 groups. As input to the clustering algorithm, we used the high-dimensional vector of escape scores encoding each trajectory (one value for every 0.2 years in the sliding window over time).

### Analyzing escape scores

The COV CATNAP tool [66] was very valuable for exploring the sea of literature on SARS-CoV-2 antibodies, and helped us identify studies with neutralization data relevant to our work. The CoVariants website (https://covariants.org/) was very useful for identifying mutations in variants of concern [26].

### Estimating fitness effects from nature

We computed fitness effects using the computational pipeline at https://github.com/neherlab/SARS2-mut-fitness-v2, which is described in Haddox et al. [45]. We ran this pipeline using a custom configuration file. See https://github.com/matsengrp/ncov-ab-escape for the custom configuration file that we used and for files reporting the estimated fitness effects.

## Acknowledgements

We thank John Huddleston and Richard Neher for useful discussions. We thank Fanchong Jian and Yunlong Cao for providing detailed answers to questions about their data. We thank Luca Sesta for providing detailed answers to questions about the pipeline for computing fitness effects.

## References

[1] Yunlong Cao, Jing Wang, Fanchong Jian, Tianhe Xiao, Weiliang Song, Ayijiang Yisimayi, Weijin Huang, Qianqian Li, Peng Wang, Ran An, et al. Omicron escapes the majority of existing SARS-CoV-2 neutralizing antibodies. Nature, 602(7898):657–663, 2022.

[2] Yunlong Cao, Ayijiang Yisimayi, Fanchong Jian, Weiliang Song, Tianhe Xiao, Lei Wang, Shuo Du, Jing Wang, Qianqian Li, Xiaosu Chen, et al. BA.2.12.1, BA.4 and BA.5 escape antibodies elicited by Omicron infection. Nature, 608(7923):593–602, 2022.

[3] Yunlong Cao, Fanchong Jian, Jing Wang, Yuanling Yu, Weiliang Song, Ayijiang Yisimayi, Jing Wang, Ran An, Xiaosu Chen, Na Zhang, et al. Imprinted SARS-CoV-2 humoral immunity induces convergent Omicron RBD evolution. Nature, 614(7948):521–529, 2023.

[4] Kang Wang, Zijing Jia, Linilin Bao, Lei Wang, Lei Cao, Hang Chi, Yaling Hu, Qianqian Li, Yunjiao Zhou, Yinan Jiang, et al. Memory B cell repertoire from triple vaccinees against diverse SARS-CoV-2 variants. Nature, 603(7903):919–925, 2022.

[5] Sarah A Clark, Lars E Clark, Junhua Pan, Adrian Coscia, Lindsay GA McKay, Sundaresh Shankar, Rebecca I Johnson, Vesna Brusic, Manish C Choudhary, James Regan, et al. SARS-CoV-2 evolution in an immunocompromised host reveals shared neutralization escape mechanisms. Cell, 184(10):2605–2617, 2021.

[6] Bryan E Jones, Patricia L Brown-Augsburger, Kizzmekia S Corbett, Kathryn Westendorf, Julian Davies, Thomas P Cujec, Christopher M Wiethoff, Jamie L Blackbourne, Beverly A Heinz, Denisa Foster, et al. The neutralizing antibody, LY-CoV555, protects against SARS-CoV-2 infection in nonhuman primates. Science Translational Medicine, 13(593):eabf1906, 2021.

[7] Johanna Hansen, Alina Baum, Kristen E Pascal, Vincenzo Russo, Stephanie Giordano, Elzbieta Wloga, Benjamin O Fulton, Ying Yan, Katrina Koon, Krunal Patel, et al. Studies in humanized mice and convalescent humans yield a SARS-CoV-2 antibody cocktail. Science, 369(6506):1010–1014, 2020.

[8] Seth J Zost, Pavlo Gilchuk, Rita E Chen, James Brett Case, Joseph X Reidy, Andrew Trivette, Rachel S Nargi, Rachel E Sutton, Naveenchandra Suryadevara, Elaine C Chen, et al. Rapid isolation and profiling of a diverse panel of human monoclonal antibodies targeting the SARS-CoV-2 spike protein. Nature Medicine, 26(9):1422–1427, 2020.

[9] Davide F Robbiani, Christian Gaebler, Frauke Muecksch, Julio CC Lorenzi, Zijun Wang, Alice Cho, Marianna Agudelo, Christopher O Barnes, Anna Gazumyan, Shlomo Finkin, et al. Convergent antibody responses to SARS-CoV-2 in convalescent individuals. Nature, 584(7821):437–442, 2020.

[10] Tyler N Starr, Nadine Czudnochowski, Zhuoming Liu, Fabrizia Zatta, Young-Jun Park, Amin Addetia, Dora Pinto, Martina Beltramello, Patrick Hernandez, Allison J Greaney, et al. SARS-CoV-2 RBD antibodies that maximize breadth and resistance to escape. Nature, 597(7874):97–102, 2021.

[11] Young-Jun Park, Dora Pinto, Alexandra C Walls, Zhuoming Liu, Anna De Marco, Fabio Benigni, Fabrizia Zatta, Chiara Silacci-Fregni, Jessica Bassi, Kaitlin R Sprouse, et al. Imprinted antibody responses against SARS-CoV-2 Omicron sublineages. Science, 378(6620):619–627, 2022.

[12] Young-Jun Park, Anna De Marco, Tyler N Starr, Zhuoming Liu, Dora Pinto, Alexandra C Walls, Fabrizia Zatta, Samantha K Zepeda, John E Bowen, Kaitlin R Sprouse, et al. Antibody-mediated broad sarbecovirus neutralization through ACE2 molecular mimicry. Science, 375(6579):449–454, 2022.

[13] Elisabetta Cameroni, John E Bowen, Laura E Rosen, Christian Saliba, Samantha K Zepeda, Katja Culap, Dora Pinto, Laura A VanBlargan, Anna De Marco, Julia di Iulio, et al. Broadly neutralizing antibodies overcome SARS-CoV-2 Omicron antigenic shift. Nature, 602(7898):664–670, 2022.

[14] Pengfei Wang, Ryan G Casner, Manoj S Nair, Jian Yu, Yicheng Guo, Maple Wang, Jasper F-W Chan, Gabriele Cerutti, Sho Iketani, Lihong Liu, et al. A monoclonal antibody that neutralizes SARS-CoV-2 variants, SARS-CoV, and other sarbecoviruses. Emerging Microbes & Infections, 11 (1):147–157, 2022.

[15] Jamie Guenthoer, Michelle Lilly, Tyler N Starr, Bernadeta Dadonaite, Klaus N Lovendahl, Jacob T Croft, Caitlin I Stoddard, Vrasha Chohan, Shilei Ding, Felicitas Ruiz, et al. Identification of broad, potent antibodies to functionally constrained regions of SARS-CoV-2 spike following a break-through infection. Proceedings of the National Academy of Sciences, 120 (23):e2220948120, 2023.

[16] Qingwen He, Lili Wu, Zepeng Xu, Xiaoyun Wang, Yufeng Xie, Yan Chai, Anqi Zheng, Jianjie Zhou, Shitong Qiao, Min Huang, et al. An updated atlas of antibody evasion by SARS-CoV-2 Omicron sub-variants including BQ.1.1 and XBB. Cell Reports Medicine, 4(4), 2023.

[17] Fanchong Jian, Jing Wang, Ayijiang Yisimayi, Weiliang Song, Yanli Xu, Xiaosu Chen, Xiao Niu, Sijie Yang, Yuanling Yu, Peng Wang, et al. Evolving antibody response to SARS-CoV-2 antigenic shift from XBB to JN.1. bioRxiv, pages 2024–04, 2024.

[18] Dami A Collier, Anna De Marco, Isabella ATM Ferreira, Bo Meng, Rawlings P Datir, Alexandra C Walls, Steven A Kemp, Jessica Bassi, Dora Pinto, Chiara Silacci-Fregni, et al. Sensitivity of SARS-CoV-2 B.1.1.7 to mRNA vaccine-elicited antibodies. Nature, 593(7857):136–141, 2021.

[19] Matthew McCallum, Alexandra C Walls, Kaitlin R Sprouse, John E Bowen, Laura E Rosen, Ha V Dang, Anna De Marco, Nicholas Franko, Sasha W Tilles, Jennifer Logue, et al. Molecular basis of immune evasion by the Delta and Kappa SARS-CoV-2 variants. Science, 374(6575):1621–1626, 2021.

[20] Amin Addetia, Luca Piccoli, James Brett Case, Young-Jun Park, Martina Beltramello, Barbara Guarino, Ha Dang, Guilherme Dias de Melo, Dora Pinto, Kaitlin Sprouse, et al. Neutralization, effector function and immune imprinting of Omicron variants. Nature, 621(7979):592–601, 2023.

[21] Delphine Planas, Nell Saunders, Piet Maes, Florence Guivel-Benhassine, Cyril Planchais, Julian Buchrieser, William-Henry Bolland, Françoise Porrot, Isabelle Staropoli, Frederic Lemoine, et al. Considerable escape of SARS-CoV-2 Omicron to antibody neutralization. Nature, 602(7898):671–675, 2022.

[22] Lihong Liu, Sho Iketani, Yicheng Guo, Jasper F-W Chan, Maple Wang, Liyuan Liu, Yang Luo, Hin Chu, Yiming Huang, Manoj S Nair, et al. Striking antibody evasion manifested by the Omicron variant of SARS-CoV-2. Nature, 602(7898):676–681, 2022.

[23] Chaoran Chen, Sarah Nadeau, Michael Yared, Philippe Voinov, Ning Xie, Cornelius Roemer, and Tanja Stadler. Cov-spectrum: analysis of globally shared SARS-CoV-2 data to identify and characterize new variants. Bioinformatics, 38(6):1735–1737, 2022.

[24] Ivan Aksamentov, Cornelius Roemer, Emma B. Hodcroft, and Richard A. Neher. Nextclade: clade assignment, mutation calling and quality control for viral genomes. Journal of Open Source Software, 6(67):3773, 2021. doi: 10.21105/joss.03773. URL 10.21105/joss.03773.

[25] Andrew Rambaut, Edward C Holmes, Áine O’Toole, Verity Hill, John T McCrone, Christopher Ruis, Louis Du Plessis, and Oliver G Pybus. A dynamic nomenclature proposal for SARS-CoV-2 lineages to assist genomic epidemiology. Nature Microbiology, 5(11):1403–1407, 2020.

[26] Emma B Hodcroft. Covariants: SARS-CoV-2 mutations and variants of interest. 2021. URL https://covariants.org/.

[27] Allison J Greaney, Tyler N Starr, Pavlo Gilchuk, Seth J Zost, Elad Binshtein, Andrea N Loes, Sarah K Hilton, John Huddleston, Rachel Eguia, Katharine HD Crawford, et al. Complete mapping of mutations to the SARS-CoV-2 spike receptor-binding domain that escape antibody recognition. Cell Host & Microbe, 29(1):44–57, 2021.

[28] Allison J Greaney, Andrea N Loes, Katharine HD Crawford, Tyler N Starr, Keara D Malone, Helen Y Chu, and Jesse D Bloom. Comprehensive mapping of mutations in the SARS-CoV-2 receptor-binding domain that affect recognition by polyclonal human plasma antibodies. Cell Host & Microbe, 29(3):463–476, 2021.

[29] Tyler N Starr, Allison J Greaney, Adam S Dingens, and Jesse D Bloom. Complete map of SARS-CoV-2 RBD mutations that escape the monoclonal antibody LY-CoV555 and its cocktail with LY-CoV016. Cell Reports Medicine, 2(4), 2021.

[30] Bernadeta Dadonaite, Katharine HD Crawford, Caelan E Radford, Ariana G Farrell, C Yu Timothy, William W Hannon, Panpan Zhou, Raiees Andrabi, Dennis R Burton, Lihong Liu, et al. A pseudovirus system enables deep mutational scanning of the full SARS-CoV-2 spike. Cell, 186 (6):1263–1278, 2023.

[31] Ruoke Wang, Qi Zhang, Jiwan Ge, Wenlin Ren, Rui Zhang, Jun Lan, Bin Ju, Bin Su, Fengting Yu, Peng Chen, et al. Analysis of SARS-CoV-2 variant mutations reveals neutralization escape mechanisms and the ability to use ACE2 receptors from additional species. Immunity, 54(7):1611–1621, 2021.

[32] Jingwen Ai, Xun Wang, Xinyi He, Xiaoyu Zhao, Yi Zhang, Yuchao Jiang, Minghui Li, Yuchen Cui, Yanjia Chen, Rui Qiao, et al. Antibody evasion of SARS-CoV-2 Omicron BA.1, BA.1.1, BA.2, and BA.3 sub-lineages. Cell Host & Microbe, 30(8):1077–1083, 2022.

[33] MacGregor Cox, Thomas P Peacock, William T Harvey, Joseph Hughes, Derek W Wright, Brian J Willett, Emma Thomson, Ravindra K Gupta, Sharon J Peacock, David L Robertson, et al. SARS-CoV-2 variant evasion of monoclonal antibodies based on in vitro studies. Nature Reviews Microbiology, 21(2):112–124, 2023.

[34] Luca Piccoli, Young-Jun Park, M Alejandra Tortorici, Nadine Czudnochowski, Alexandra C Walls, Martina Beltramello, Chiara Silacci-Fregni, Dora Pinto, Laura E Rosen, John E Bowen, et al. Mapping neutralizing and immunodominant sites on the SARS-CoV-2 spike receptor-binding domain by structure-guided high-resolution serology. Cell, 183(4):1024–1042, 2020.

[35] James Hadfield, Colin Megill, Sidney M Bell, John Huddleston, Barney Potter, Charlton Callender, Pavel Sagulenko, Trevor Bedford, and Richard A Neher. Nextstrain: real-time tracking of pathogen evolution. Bioinformatics, 34(23):4121–4123, 2018.

[36] Allison J Greaney, Tyler N Starr, and Jesse D Bloom. An antibody-escape estimator for mutations to the SARS-CoV-2 receptor-binding domain. Virus Evolution, 8(1):veac021, 2022.

[37] Ayijiang Yisimayi, Weiliang Song, Jing Wang, Fanchong Jian, Yuanling Yu, Xiaosu Chen, Yanli Xu, Sijie Yang, Xiao Niu, Tianhe Xiao, et al. Repeated Omicron exposures override ancestral SARS-CoV-2 immune imprinting. Nature, 625(7993):148–156, 2024.

[38] Yatish Turakhia, Bryan Thornlow, Angie Hinrichs, Jakob McBroome, Nicolas Ayala, Cheng Ye, Kyle Smith, Nicola De Maio, David Haussler, Robert Lanfear, et al. Pandemic-scale phylogenomics reveals the SARS-CoV-2 recombination landscape. Nature, 609(7929):994–997, 2022.

[39] Tomokazu Tamura, Jumpei Ito, Keiya Uriu, Jiri Zahradnik, Izumi Kida, Yuki Anraku, Hesham Nasser, Maya Shofa, Yoshitaka Oda, Spyros Lytras, et al. Virological characteristics of the SARS-CoV-2 XBB variant derived from recombination of two Omicron subvariants. Nature Communications, 14(1):2800, 2023.

[40] James F Crow and Motoo Kimura. Evolution in sexual and asexual populations. The American Naturalist, 99(909):439–450, 1965.

[41] Matthew Hegreness, Noam Shoresh, Daniel Hartl, and Roy Kishony. An equivalence principle for the incorporation of favorable mutations in asexual populations. Science, 311(5767):1615–1617, 2006.

[42] Katy C Kao and Gavin Sherlock. Molecular characterization of clonal interference during adaptive evolution in asexual populations of saccharomyces cerevisiae. Nature Genetics, 40(12):1499–1504, 2008.

[43] Gregory I Lang, Daniel P Rice, Mark J Hickman, Erica Sodergren, George M Weinstock, David Botstein, and Michael M Desai. Pervasive genetic hitchhiking and clonal interference in forty evolving yeast populations. Nature, 500(7464):571–574, 2013.

[44] Jesse D Bloom and Richard A Neher. Fitness effects of mutations to SARS-CoV-2 proteins. Virus Evolution, 9(2):vead055, 2023.

[45] Hugh K Haddox, Georg Angehrn, Luca Sesta, Chris Jennings-Shaffer, Seth D Temple, Jared G Galloway, William S DeWitt, Jesse D Bloom, Frederick A Matsen, and Richard A Neher. The mutation rate of SARS-CoV-2 is highly variable between sites and is influenced by sequence context, genomic region, and RNA structure. bioRxiv, 2025.

[46] Tyler N Starr, Allison J Greaney, Cameron M Stewart, Alexandra C Walls, William W Hannon, David Veesler, and Jesse D Bloom. Deep mutational scans for ACE2 binding, RBD expression, and antibody escape in the SARS-CoV-2 Omicron BA.1 and BA.2 receptor-binding domains. PLoS Pathogens, 18(11):e1010951, 2022.

[47] Leander Witte, Viren A Baharani, Fabian Schmidt, Zijun Wang, Alice Cho, Raphael Raspe, Camila Guzman-Cardozo, Frauke Muecksch, Marie Canis, Debby J Park, et al. Epistasis lowers the genetic barrier to SARS-CoV-2 neutralizing antibody escape. Nature Communications, 14(1):302, 2023.

[48] Tyler N Starr, Allison J Greaney, William W Hannon, Andrea N Loes, Kevin Hauser, Josh R Dillen, Elena Ferri, Ariana Ghez Farrell, Bernadeta Dadonaite, Matthew McCallum, et al. Shifting mutational constraints in the SARS-CoV-2 receptor-binding domain during viral evolution. Science, 377(6604):420–424, 2022.

[49] Alief Moulana, Thomas Dupic, Angela M Phillips, Jeffrey Chang, Serafina Nieves, Anne A Roffler, Allison J Greaney, Tyler N Starr, Jesse D Bloom, and Michael M Desai. Compensatory epistasis maintains ACE2 affinity in SARS-CoV-2 Omicron BA.1. Nature Communications, 13(1):7011, 2022.

[50] Ping He, Banghui Liu, Xijie Gao, Qihong Yan, Rongjuan Pei, Jing Sun, Qiuluan Chen, Ruitian Hou, Zimu Li, Yanjun Zhang, et al. SARS-CoV-2 Delta and Omicron variants evade population antibody response by mutations in a single spike epitope. Nature Microbiology, 7(10):1635–1649, 2022.

[51] Meng Yuan, Xueyong Zhu, Wan-ting He, Panpan Zhou, Chengzi I Kaku, Tazio Capozzola, Connie Y Zhu, Xinye Yu, Hejun Liu, Wenli Yu, et al. A broad and potent neutralization epitope in SARS-related coronaviruses. Proceedings of the National Academy of Sciences, 119(29):e2205784119, 2022.

[52] Min Huang, Lili Wu, Anqi Zheng, Yufeng Xie, Qingwen He, Xiaoyu Rong, Pu Han, Pei Du, Pengcheng Han, Zengyuan Zhang, et al. Atlas of currently available human neutralizing antibodies against SARS-CoV-2 and escape by Omicron sub-variants BA.1/BA.1.1/BA.2/BA.3. Immunity, 55(8):1501–1514, 2022.

[53] Rosario Miralles, Philip J Gerrish, Andrés Moya, and Santiago F Elena. Clonal interference and the evolution of RNA viruses. Science, 285(5434):1745–1747, 1999.

[54] Natalja Strelkowa and Michael Lässig. Clonal interference in the evolution of influenza. Genetics, 192(2):671–682, 2012.

[55] Katherine S Xue, Terry Stevens-Ayers, Angela P Campbell, Janet A Englund, Steven A Pergam, Michael Boeckh, and Jesse D Bloom. Parallel evolution of influenza across multiple spatiotemporal scales. Elife, 6:e26875, 2017.

[56] Frances C Welsh, Rachel T Eguia, Juhye M Lee, Hugh K Haddox, Jared Galloway, Nguyen Van Vinh Chau, Andrea N Loes, John Huddleston, C Yu Timothy, Mai Quynh Le, et al. Age-dependent heterogeneity in the antigenic effects of mutations to influenza hemagglutinin. Cell Host & Microbe, 32(8):1397–1411, 2024.

[57] Caleb R Carr, Katharine HD Crawford, Michael Murphy, Jared G Galloway, Hugh K Haddox, Frederick A Matsen, Kristian G Andersen, Neil P King, and Jesse D Bloom. Deep mutational scanning reveals functional constraints and antibody-escape potential of Lassa virus glycoprotein complex. Immunity, 57(9):2061–2076, 2024.

[58] Bernadeta Dadonaite, Jack Brown, Teagan E McMahon, Ariana G Farrell, Marlin D Figgins, Daniel Asarnow, Cameron Stewart, Jimin Lee, Jenni Logue, Trevor Bedford, et al. Spike deep mutational scanning helps predict success of SARS-CoV-2 clades. Nature, 631(8021):617–626, 2024.

[59] Ruipeng Lei, Enya Qing, Abby Odle, Meng Yuan, Chaminda D Gunawardene, Timothy JC Tan, Natalie So, Wenhao O Ouyang, Ian A Wilson, Tom Gallagher, et al. Functional and antigenic characterization of SARS-CoV-2 spike fusion peptide by deep mutational scanning. Nature Communications, 15(1):4056, 2024.

[60] Mellissa C Alcantara, Yusuke Higuchi, Yuhei Kirita, Satoaki Matoba, and Atsushi Hoshino. Deep mutational scanning to predict escape from bebtelovimab in SARS-CoV-2 Omicron subvariants. Vaccines, 11(3):711, 2023.

[61] Bernadeta Dadonaite, Jenny J Ahn, Jordan T Ort, Jin Yu, Colleen Furey, Annie Dosey, William W Hannon, Amy L Vincent Baker, Richard J Webby, Neil P King, et al. Deep mutational scanning of H5 hemagglutinin to inform influenza virus surveillance. PLoS Biology, 22(11):e3002916, 2024.

[62] Brendan B Larsen, Teagan McMahon, Jack T Brown, Zhaoqian Wang, Caelan E Radford, James E Crowe Jr, David Veesler, and Jesse D Bloom. Functional and antigenic landscape of the Nipah virus receptor binding protein. bioRxiv, 2024.

[63] Arjun K Aditham, Caelan E Radford, Caleb R Carr, Naveen Jasti, Neil P King, and Jesse D Bloom. Deep mutational scanning of Rabies glycoprotein defines mutational constraint and antibody-escape mutations. bioRxiv, pages 2024–12, 2024.

[64] Caroline Kikawa, Catiana H Cartwright-Acar, Jackson B Stuart, Maya Contreras, Lisa M Levoir, Matthew J Evans, Jesse D Bloom, and Leslie Goo. The effect of single mutations in Zika virus envelope on escape from broadly neutralizing antibodies. Journal of Virology, 97(11):e01414–23, 2023.

[65] Caelan E Radford, Philipp Schommers, Lutz Gieselmann, Katharine HD Crawford, Bernadeta Dadonaite, C Yu Timothy, Adam S Dingens, Julie Overbaugh, Florian Klein, and Jesse D Bloom. Mapping the neutralizing specificity of human anti-HIV serum by deep mutational scanning. Cell Host & Microbe, 31(7):1200–1215, 2023.

[66] Hyejin Yoon, Jennifer Macke, Anthony P West Jr, Brian Foley, Pamela J Bjorkman, Bette Korber, and Karina Yusim. CATNAP: a tool to compile, analyze and tally neutralizing antibody panels. Nucleic Acids Research, 43 (W1):W213–W219, 2015.

