## Supplementary Information for "Clonal interference and changing selective pressures shape the escape of SARS-CoV-2 from hundreds of antibodies"

January 22, 2025

### Figures and Tables

### References

[1] Yunlong Cao, Fanchong Jian, Jing Wang, Yuanling Yu, Weiliang Song, Ayijiang Yisimayi, Jing Wang, Ran An, Xiaosu Chen, Na Zhang, et al. Imprinted SARS-CoV-2 humoral immunity induces convergent Omicron RBD evolution. *Nature*, 614(7948):521–529, 2023.

[2] Tyler N Starr, Allison J Greaney, William W Hannon, Andrea N Loes, Kevin Hauser, Josh R Dillen, Elena Ferri, Ariana Ghez Farrell, Bernadeta Dadonaite, Matthew McCallum, et al. Shifting mutational constraints in the SARS-CoV-2 receptor-binding domain during viral evolution. *Science*, 377(6604):420–424, 2022.

[3] Tyler N Starr, Allison J Greaney, Cameron M Stewart, Alexandra C Walls, William W Hannon, David Veesler, and Jesse D Bloom. Deep mutational scans for ACE2 binding, RBD expression, and antibody escape in the SARS-CoV-2 Omicron BA.1 and BA.2 receptor-binding domains. *PLoS Pathogens*, 18(11):e1010951, 2022.

| source | number of antibodies |
|---|---|
| WT vaccinees | 205 |
| WT convalescents | 321 |
| BA.1 convalescents | 462 |
| BA.2 convalescents | 470 |
| BA.5 convalescents | 145 |

**Table S1: Number of antibodies per source.** Cao et al. isolated antibodies from humans with a variety of exposure histories. For the 1,603 antibodies from our analysis, the table shows how many antibodies were isolated from a human with a given exposure history.

tab:sources

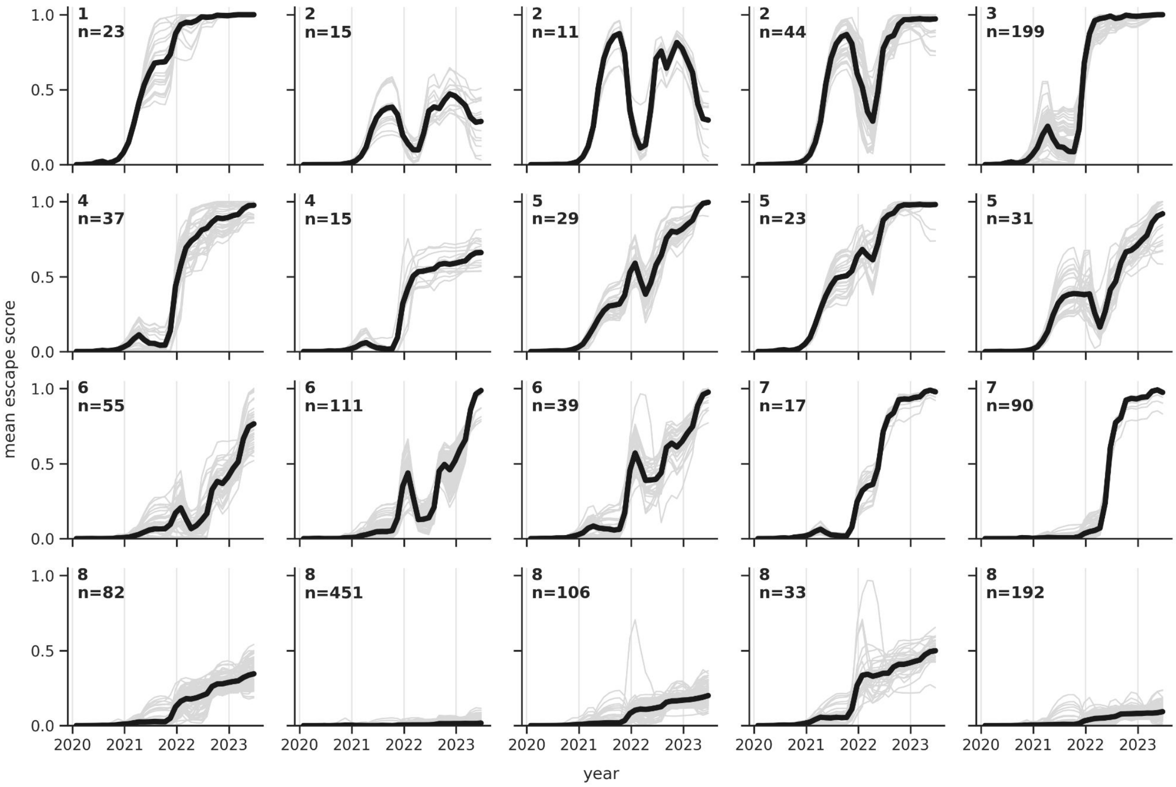


**Figure S1: The 20 clusters of escape trajectories generated by $k$-means clustering.** Each panel shows data for a different cluster. In each panel, thin gray lines show trajectories for individual antibodies within a cluster, while the bold black line shows the average trajectory. The title of each panel indicates the consolidated cluster number 1-8 from Figure 2B (e.g., the three panels with the title of "2" were all combined into a single cluster) as well as the number of antibodies per panel. fig:kmeans

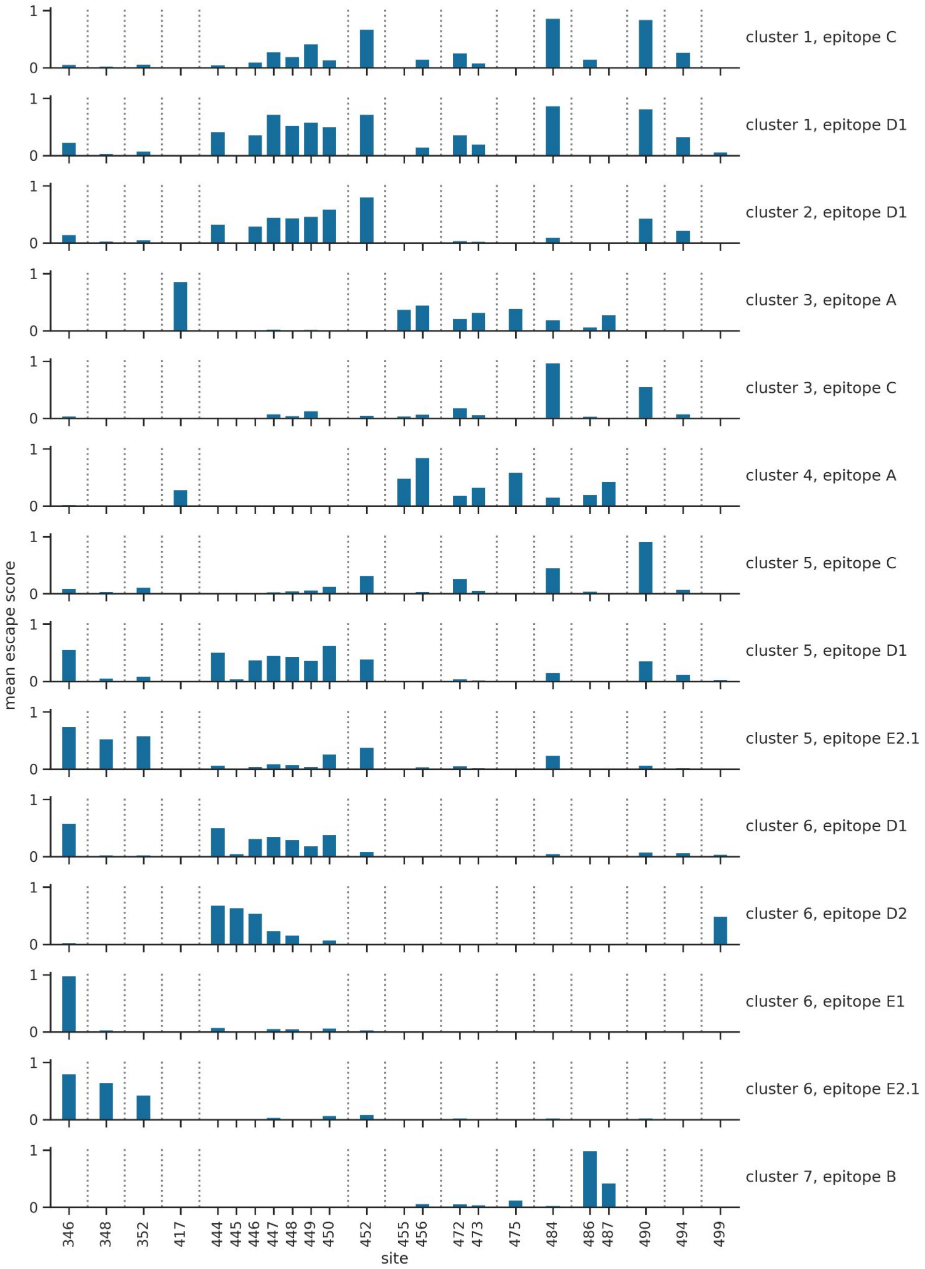


**Figure S2: Average site-level escape scores of all antibodies in a given escape cluster and epitope.** Escape clusters come from this paper, while epitope clusters come from Cao et al. [1]. The plot only shows data for sites where the average site-level escape score is >0.3 for at least one group of antibodies. fig:dms˙profiles

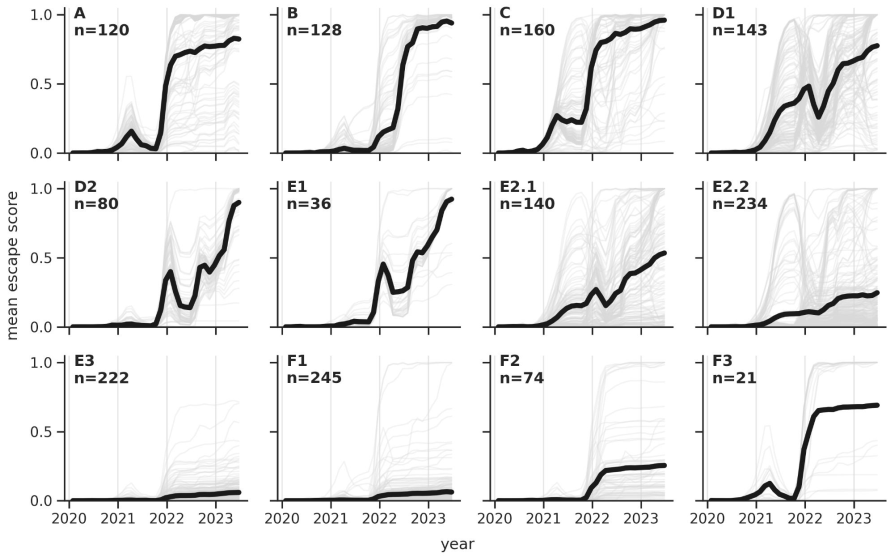


**Figure S3: Escape trajectories clustered by antibody epitopes.** Each panel shows data for a different epitope cluster, as defined by Cao et al. [1]. In each panel, thin gray lines show trajectories for individual antibodies within an epitope cluster, while the bold black line shows the average trajectory. The title of each panel indicates the epitope as well as the number of antibodies per panel. fig:trajectories˙by˙epitope

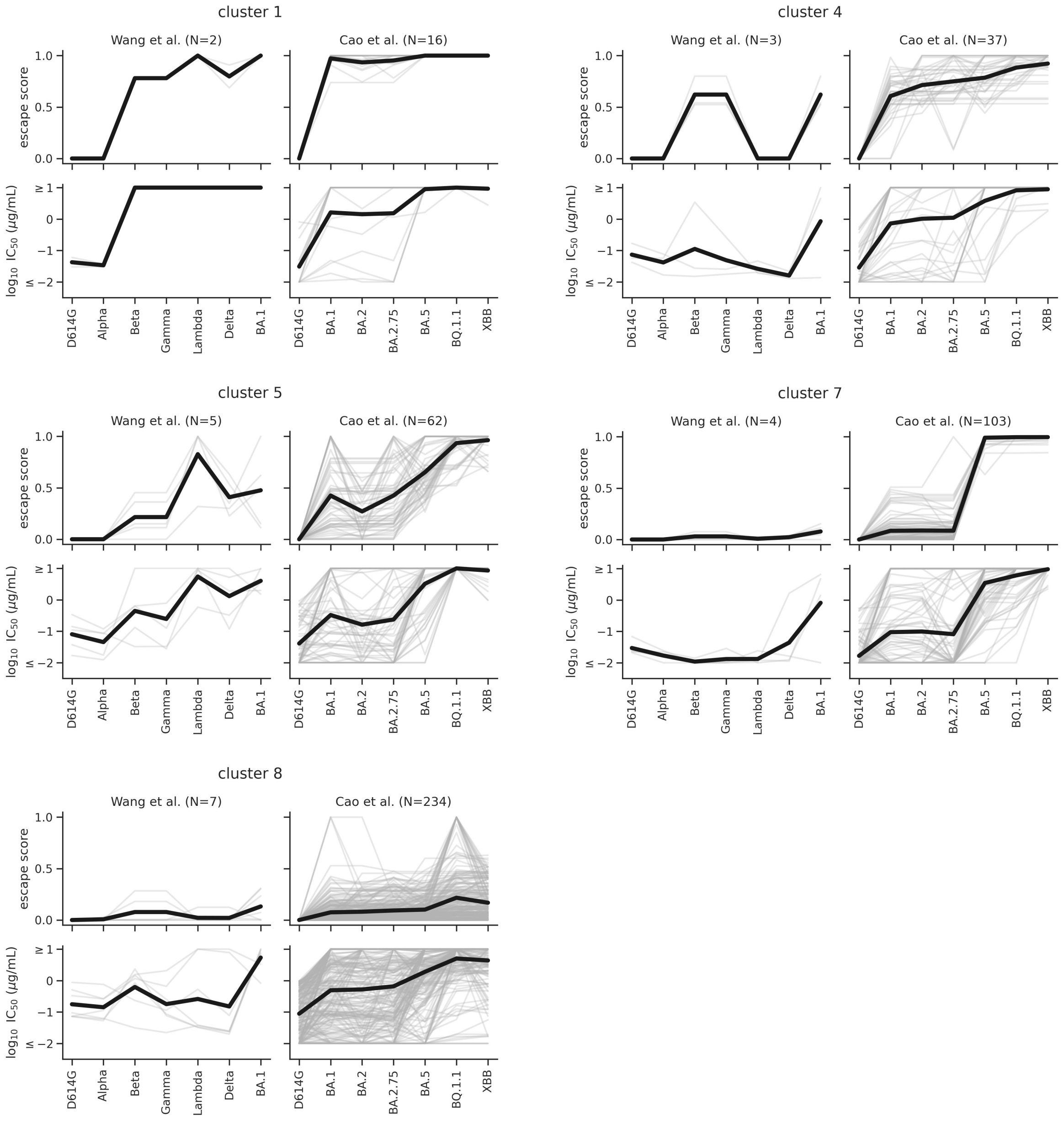


**Figure S4: Comparing predicted escape scores and pseudovirus neutralization data for the remaining clusters.** This figure is similar to Figure 4B-E, but shows data for the remaining clusters.

fig:validation`supp

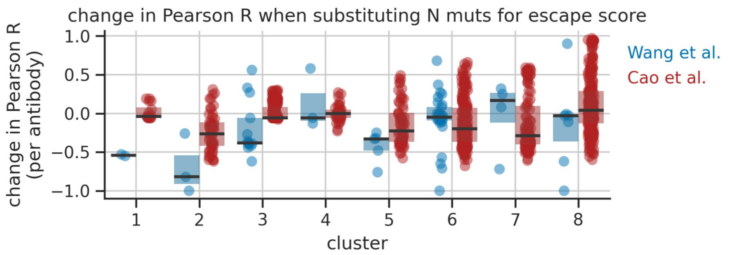


**Figure S5: Escape scores capture more than sequence divergence.** This figure is similar to Figure 4B, but shows the change in per-antibody correlation coefficients when they are computed using the number of RBD mutations in a spike variant rather than its predicted escape score for that antibody. Negative changes mean that the correlation coefficients are lower when using the number of RBD mutations. We impose a floor of $-1$ on these values. For clusters 1-3 and 5-7, the median correlation coefficient per cluster is substantially less than zero for at least one study. Note: in cases where the median change is close to zero, that does not necessarily indicate that the escape scores for that group of antibodies are incorrect, it merely indicates the pattern aligns with sequence divergence. fig:validation˙delta˙corr

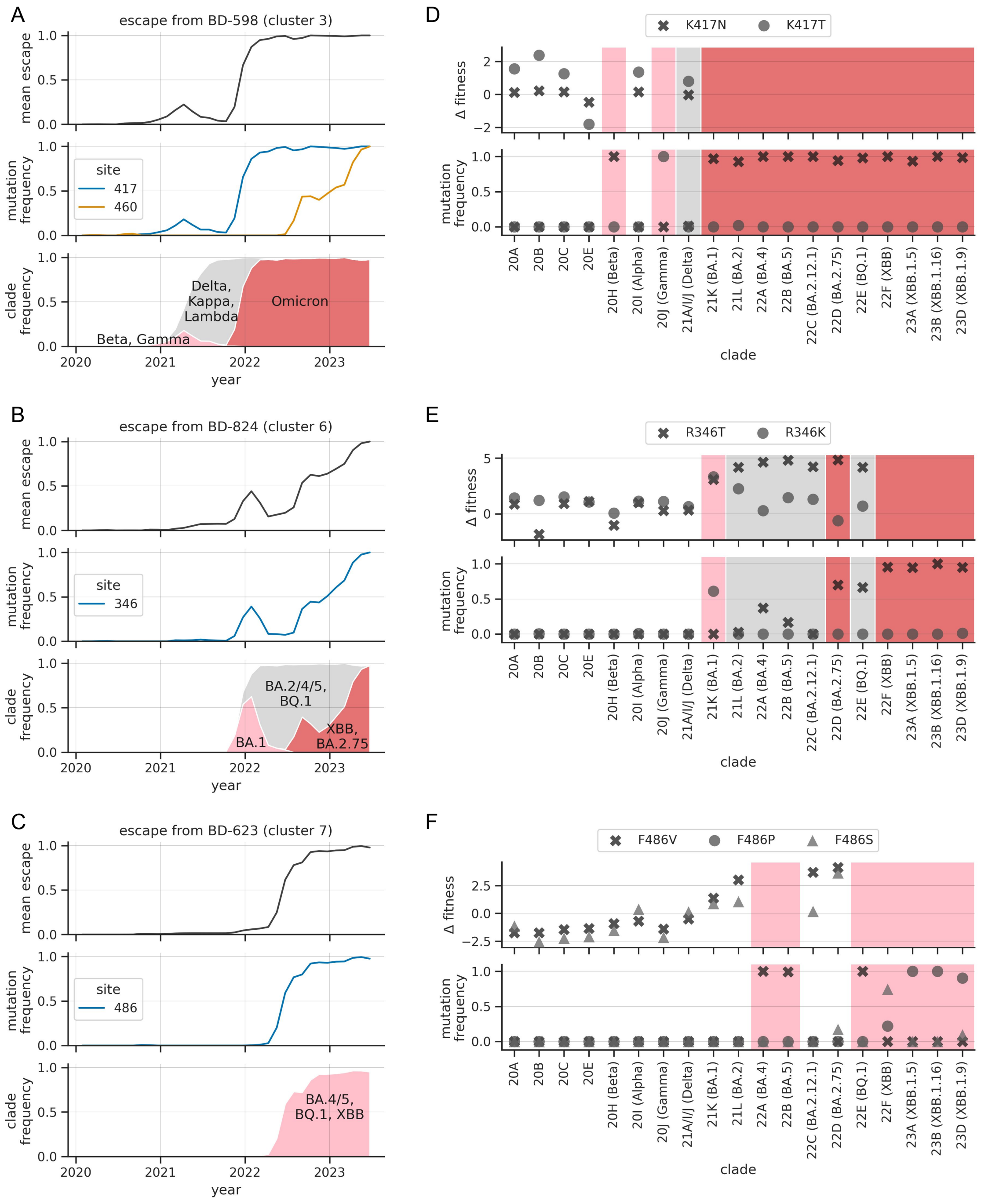


**Figure S6:** Same as Figure 5, but panels A-C show data for a different set of example antibodies and panels D-F show data for escape mutations associated with those antibodies.

fig:clade˙displacement˙supp

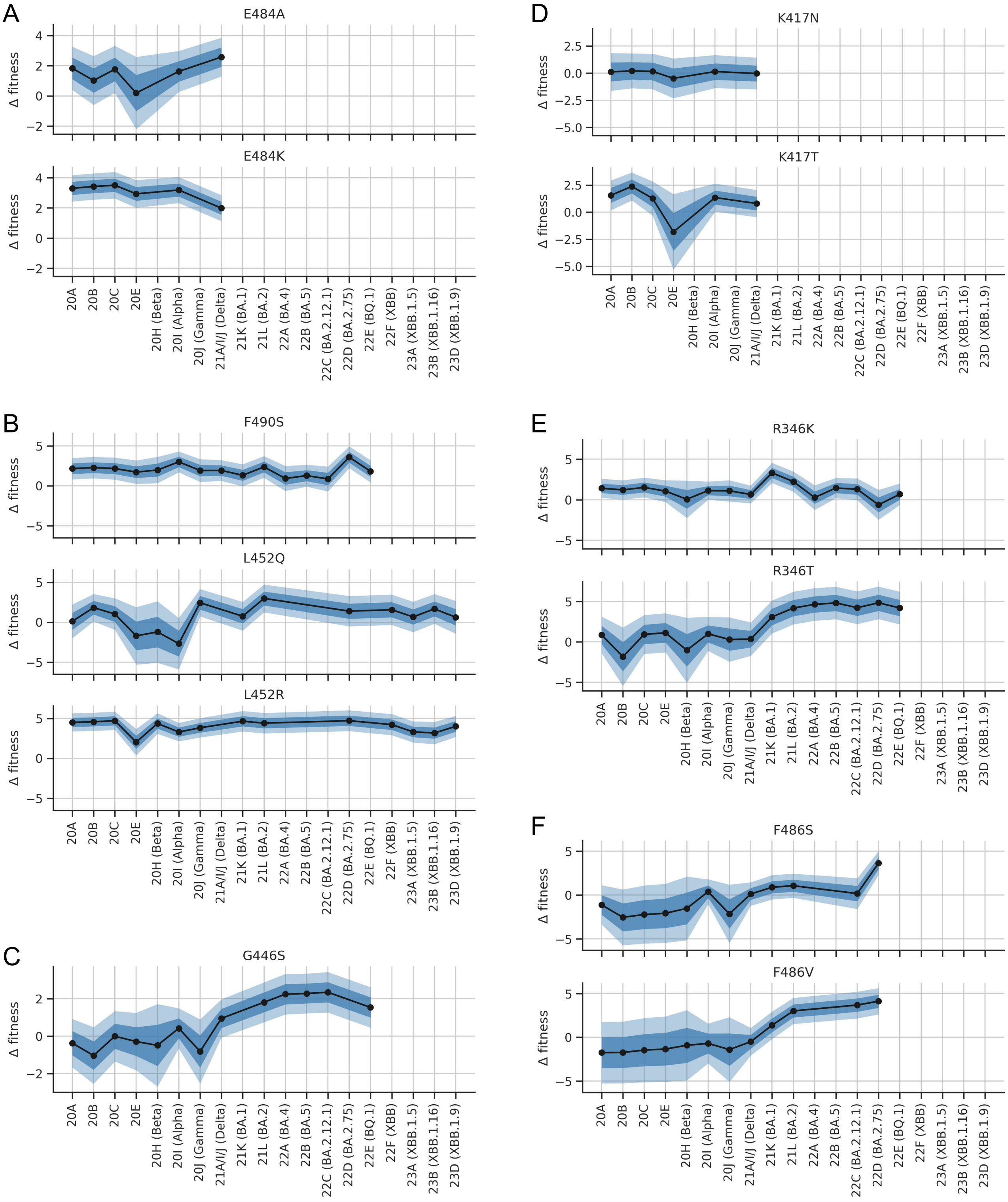


**Figure S7: Levels of uncertainty in estimated fitness effects.** Each panel shows fitness effects for a given group of mutations. Panels A-C show groups of mutations from Figure 5D-F, while panels D-F show groups of mutations from Fig. S6D-F. In each panel, each subplot shows data for a given mutation. For a given clade on the x-axis, the black dots show a mutation's fitness effect, computed as the mean of the posterior distribution over fitness effects, while the dark blue region extending above and below each dot shows the mean plus or minus the standard deviation of the posterior distribution. The light-blue region shows the mean plus or minus two times the standard deviation.

fig:fitness˙uncertainty

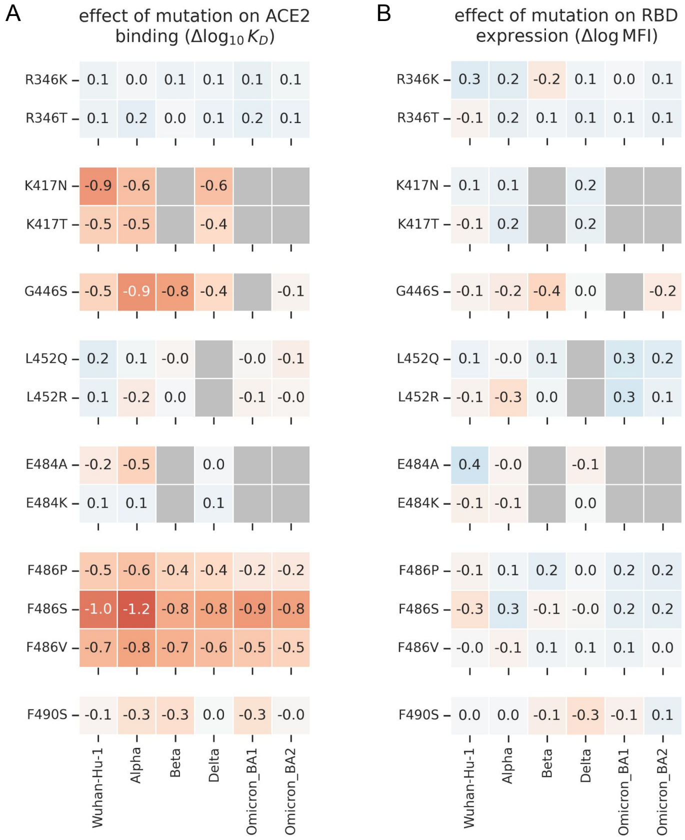


**Figure S8: Effects of mutations on RBD-ACE2 binding and RBD expression.** **(A)** Effects on binding. **(B)** Effects on expression. In both panels, the color scale ranges from $-2$ to 2, with negative numbers indicating that the mutation decreases binding or expression. These data come from two studies by Starr et al. [2, 3]. fig:rbd dms
